# Chronology-based architecture of descending circuits that underlie the development of locomotor repertoire after birth

**DOI:** 10.1101/425587

**Authors:** Avinash Pujala, Minoru Koyama

**Affiliations:** Janelia Research Campus, HHMI, Ashbum VA 20147

## Abstract

The emergence of new and increasingly sophisticated behaviors after birth is accompanied by dramatic increase of newly established synaptic connections in the nervous system. Little is known, however, of how nascent connections are organized to support such new behaviors alongside existing ones. To understand this, in the larval zebrafish we examined the development of spinal pathways from hindbrain V2a neurons and the role of these pathways in the development of locomotion. We found that new projections are continually layered laterally to existing neuropil, and give rise to distinct pathways that function in parallel to existing pathways. Across these chronologically layered pathways, the connectivity patterns and biophysical properties vary systematically to support a behavioral repertoire with a wide range of kinematics and dynamics. Such layering of new parallel circuits equipped with systematically changing properties may be central to the postnatal diversification and increasing sophistication of an animal’s behavioral repertoire.

## Introduction

At the time of birth, most animals are only capable of a limited set of reflexive behaviors that ensure their immediate survival. However, as development progresses, their behavioral repertoire rapidly diversifies and they become capable of producing increasingly refined and cognitive behaviors (Kagan and Herschkowitz 2006; Harlow and Harlow 1965; Fox 1965; Drapeau et al. 2002). Concurrent with such changes, many new connections are rapidly being formed in the nervous system (Gilmore, Knickmeyer, and Gao 2018; Semple et al. 2013; Levitt 2003), suggesting that the formation of new connections is linked to the development of the behavioral repertoire. However, despite many studies that carried out circuit-level examination of the postnatal formation of new connections (Morrie and Feller 2016; Polley et al. 2013; Stein and Stanford 2013; Kano and Watanabe 2013) very little is known still of how nascent connections are organized to support a behavioral repertoire that grows by including increasingly more sophisticated behaviors while retaining vital reflexive behaviors.

Recent studies in developing zebrafish have shed some light on the neural underpinnings of behavior development. As in most animals, the behavioral repertoire of zebrafish quickly diversifies after birth (hatching) to include more sophisticated behaviors (Drapeau et al. 2002; McLean and Fetcho 2009). At the time of birth, a zebrafish is mostly quiescent but in response to a tactile stimulus exhibits crude locomotor behavior that consists of fast and powerful left-right alternation of whole-body bending (Figure 1A, Evoked swim). During this behavior the fish’s head makes large side-to-side sweeps nearly at the same time and in the same direction as the tail (McLean and Fetcho 2009, Figure 1B). This indicates near instantaneous activation of axial muscles across the full extent of the fish’s body. However, by two days after birth, the fish also spontaneously exhibit a more refined locomotor behavior wherein only the caudal portion of the tail moves (Figure 1 A, Spontaneous swim) with relatively weaker and slower wave-like undulating movements (McLean and Fetcho 2009, Figure 1A; Mirat et al. 2013). Such movements are indicative of slow propagation of the underlying axial muscle activity from the rostral to the caudal end of the moving part of the tail. With this type of tail-restricted locomotion the fish is able to move forward by keeping its head, and therefore gaze, stable. This ability may be critical for the subsequent development of visually-guided behaviors such as prey capture. The crude and fast locomotion pattern that appears first is controlled by spinal Interneurons that are born well before the birth of a zebrafish (McLean and Fetcho 2009; Kimura, Okamura, and Higashijima 2006; Eklöf-Ljunggren et al. 2012) whereas the more refined and slower locomotion pattern that emerges after the birth of an animal is controlled by spinal Interneurons that are born around the time of birth of an animal (McLean et al. 2007; Satou, Kimura, and Higashijima 2012; McLean and Fetcho 2009). This indicates that the sequential emergence of new and more sophisticated locomotor patterns during development is mediated at the level of the spinal cord by the addition of distinct functional groups of neurons rather than by fine control of a single functional group (Fetcho and McLean 2010). Similarly in the mammalian spinal cord, it has been shown more recently that distinct functional groups emerge in sequence during development (Tripodi, Stepien, and Arber 2011). This finding has led to the idea that even in mammals, the development of the locomotor repertoire is supported by the addition of new functional groups (Tripodi and Arber 2012). However, it has yet to be revealed how the connections from the brain to emerging spinal functional groups are established so that new and existing spinal groups can be recruited appropriately by the brain to support the diversification of the locomotor repertoire.

**Figure 1.**
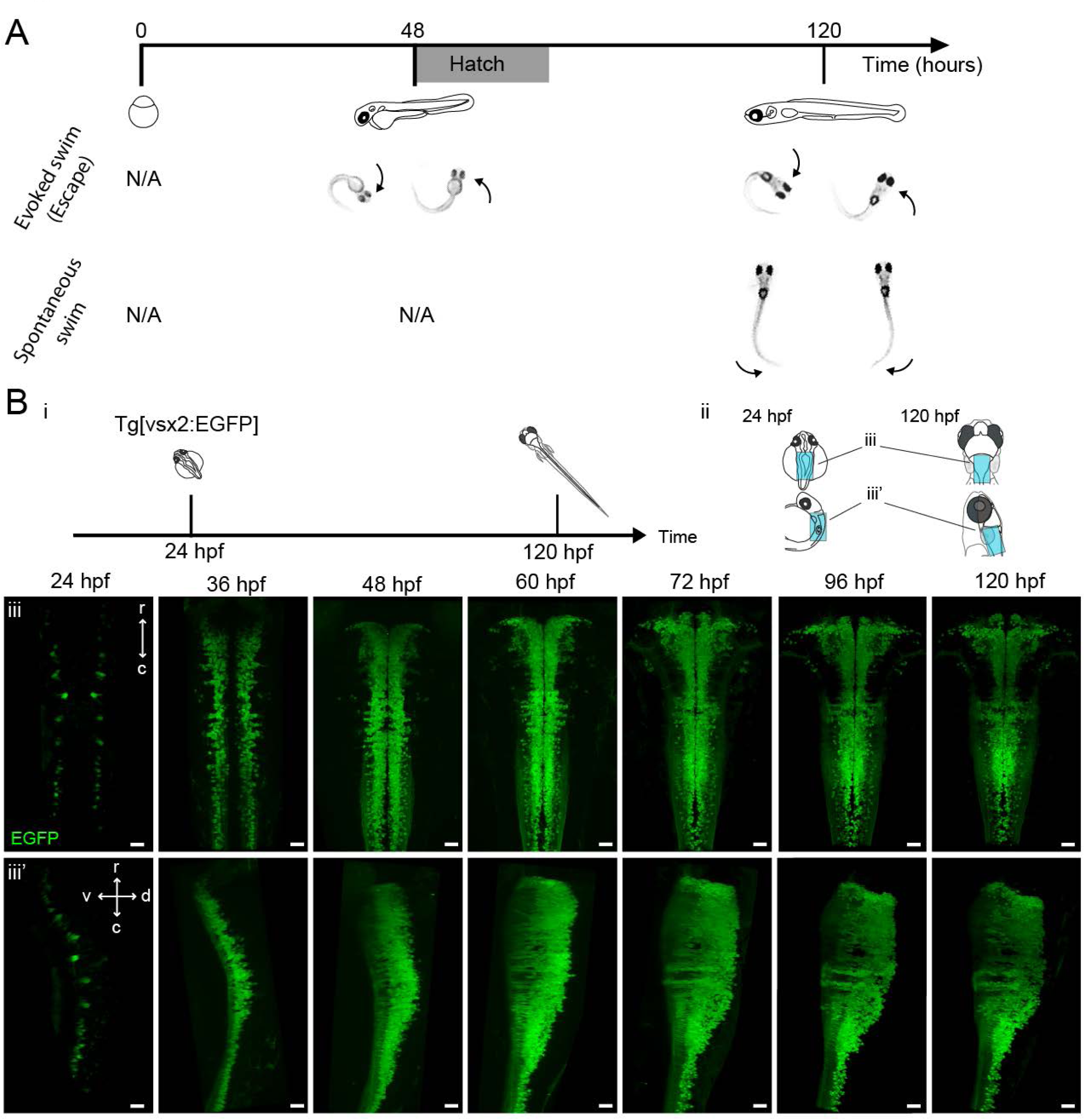
Development of zebrafish locomotor behaviors and genesis of hindbrain V2a neurons. **A.** Development of locomotor behaviors in zebrafish. Zebrafish are capable of generating strong escape swims in response to strong stimuli (Evoked swim) by the time of birth (48-72 hpf, shaded area). During this behavior, a fish’s head oscillates side-to-side as part of whole-body bending (arrows). By 120 hpf, a fish also exhibits weaker swims spontaneously (Spontaneous swim). During this behavior, the bending of the body mostly occurs in the caudal portion of the tail and the head stays mostly stable during swimming. **B.** Emergence of hindbrain V2a neurons expressing EGFP. (i) Timing of experiments. (ii) Regions displayed (cyan patch) in subsequent panels. (iii) Top-down and side views of hindbrain V2a neurons from 24 hpf to 120 hpf. r, rostral; c, caudal; d, dorsal; v, ventral. Scale bars, 30 um.

Locomotion related signals from the brain reach the spinal cord by way of excitatory reticulospinal neurons of the hindbrain (Dubuc et al. 2008; Grillner and Georgopoulos 1996; Jordan et al. 2008; Roberts et al. 2008). V2a neurons in the hindbrain, similar to the ones in the spinal cord, are glutamatergic and project their axons ipsilaterally (Cepeda-Nieto, Pfaff, and Varela-Echavarría 2005; Kinkhabwala et al. 2011; Kimura et al. 2013; Bouvier et al. 2015). They include excitatory reticulospinal neurons (Cepeda-Nieto, Pfaff, and Varela-Echavarría 2005; Kimura et al. 2013; Bouvier et al. 2015) and are capable of initiating and terminating locomotion (Kimura et al. 2013; Bouvier et al. 2015). Interestingly, the hindbrain V2a neurons in larval zebrafish show a topographical organization that is related to differentiation time such that newly-born neurons stack dorsally to pre-existing ones (Kinkhabwala et al. 2011). Furthermore, in a subset of the hindbrain V2a neurons, it has been shown that the ventral, and therefore early-born, population gets recruited during fast locomotion while the dorsal, and therefore late-born, population gets recruited during slow locomotion (Kinkhabwala et al. 2011). Even though it is unknown whether this finding holds true for the entire hindbrain V2a population, it suggests that, even in the hindbrain V2a neurons, distinct functional groups emerge in sequence and contribute to the development of locomotor behaviors. Such hindbrain organization raises the possibility that a new hindbrain functional group connects selectively to an age-matched spinal functional group to produce a novel locomotion pattern. However, it is also possible that hindbrain functional groups emerging in succession provide increasingly finer excitatory drive to all spinal groups, but each hindbrain group recruits only the age-matched spinal group because of the higher excitability of the latter compared to its predecessors (McLean et al. 2007). Indeed, in the case of midbrain descending pathways, the recruitment of spinal neurons is determined not by selectivity of the descending inputs these neurons receive but by the biophysical properties they express (Wang and McLean 2014). Thus, it remains to be resolved how the the hindbrain descending neurons establish connections to the spinal functional groups during development to support the postnatal diversification and increased sophistication of locomotor patterns.

Here, we examined the development of spinal pathways from the hindbrain V2a neurons and the role of these pathways in the development of locomotion. We show that early-born V2a descending neurons that project to the spinal cord early in development are only recruited during the stimulus-elicited crude and fast locomotion that appears early in development whereas their late-born counterparts that project to the spinal cord late in development are recruited during the more refined and slower spontaneous locomotion that appears later in development. Moreover, the spinal projections of these two populations form spatially-distinct neuropil layers and give rise to parallel pathways instead of forming a series of non-selective pathways that differ in the strengths of their connections. Furthermore, these parallel pathways differ in their connectivity patterns and biophysical properties in a manner suitable for the locomotor behavior they participate in. The early-born group makes direct connections to motoneurons and expresses biophysical properties with fast time constants. Both these features of the early-born group are well-suited for producing the crude locomotion that appears early in development in that they support the near instantaneous activation of axial muscles across the whole rostrocaudal extent of the fish’s body. On the other hand, the late-born group indirectly connects to caudal motoneurons via spinal interneurons and expresses biophysical properties with slow kinetics. These features make this group more suitable for producing the refined locomotion that appears later in development in that they support the slow rostrocaudal propagation of axial muscle activity in the caudal portion of the tail. Indeed, ablation of each group produced deficits in distinct motor patterns: ablation of the early-born group weakened the sensory-elicited crude and fast locomotion whereas ablation of the late-born group affected the more refined and slower spontaneous locomotion. Altogether, we reveal in the descending circuits a chronologically-layered parallel architecture underlying the diversification and increased sophistication of motor patterns in larval zebrafish. Even though chronotopic neuropil organization similar to what we have described here has been observed in many neural systems (Espinosa and Luo 2008; Tripodi, Stepien, and Arber 2011; Voigt, De Lima, and Beckmann 1993; Walsh and Guillery 1985; Kulkarni et al. 2016; David J. Brierley et al. 2009; D. J. Brierley et al. 2012), it has not been demonstrated in these systems how such organization relates to the postnatal development of behaviors. The findings we report here suggest that the chronological layering of parallel circuits and the systematic variation in the associated connectivity patterns and biophysical properties are fundamental attributes of the nervous system that form the basis of an animal’s ability to exhibit new and increasingly sophisticated behaviors after birth while maintaining vital reflexive behaviors.

## Results

### 1. Neuronal birth order dictates cell body position and order of spinal projections in hindbrain V2a population

To link the development of the locomotor repertoire to the development of hindbrain V2a neurons and their spinal pathways, we systematically examined the ontogeny of these neurons, their topography and the onset of their projections to the spinal cord until 120 hours post-fertilization (hpf) when larvae are capable of exhibiting both stimulus-evoked crude locomotion that appears by 48 to 72 hpf and more refined spontaneous locomotion that appears by 96 to 120 hpf (Figure 1A) (Drapeau et al. 2002; McLean and Fetcho 2009).

We first examined the ontogeny of hindbrain V2a neurons from 24 hours post fertilization (hpf) to 120 hpf with time lapse imaging (Figure 1B, n=8 for each timepoint) of a transgenic line expressing EGFP under the control of the promoter of *vsx2,* a transcription factor specific to V2a neurons. This transcription factor has been shown to become active after the final cell division (Kimura, Satou, and Higashijima 2008). Therefore, the onset of EGFP expression in a V2a neuron in our transgenic line indicates the time of differentiation - which we use synonymously with the time of birth - of this neuron. At 24 hpf, one or two pairs of V2a neurons appeared in each rhombomere (Figure 1B iii, iii’, 24 hpf), and then the number of V2a neurons increased dramatically until 60 hpf (Figure 1B iii, iii’, 60 hpf). After 72 hpf, the V2a cluster showed no drastic visible changes in the overall shape (Figure 1B iii, iii’, 72-120 hpf). As a fish is still only capable of generating crude forms of locomotion at 72 hpf, this suggests that the dramatic change in the arrangement of V2a neurons before 72 hpf may not play a direct role in the development of refined locomotion.

We then examined where V2a neurons born at different time points resided in the hindbrains of 120 hpf fish that are capable of generating both the crude and refined forms of locomotion. We did this by photoconverting Kaede, a fluorescent photoconvertible protein, which was expressed under the control of the *vsx2* promoter. We converted Kaede throughout the fish at different time points (24-96 hpf) in different groups of fish and imaged each of these groups at 120 hpf (Figure 2 supplement 1A i, n=8 for each timepoint). Based on the presence of converted Kaede (shown in magenta), we distinguished neurons that already had Kaede at the time of photoconversion from those that started to express Kaede after the time of photoconversion (Kimura, Okamura, and Higashijima 2006; Caron et al. 2008). In most rhombomeres, V2a neurons born by 24 hpf were located in the most lateral portions of the brain region occupied by this group of neurons (Figure 2 supplement 1C, 24 hpf conversion, magenta cells), whereas neurons born afterwards gradually filled up the space medial to the pre-existing ones (Figure 2 supplement 1C, 36-72 hpf conversion). Compared to photoconversion at earlier time points, photoconversion at 72 hpf revealed relatively few, if any, unconverted green cells, except in the rostral hindbrain (Figure 2 supplement 1C, 72 hpf conversion). This indicated that the neurogenesis of hindbrain V2a neurons was mostly complete by 72 hpf when fish are still only capable of producing the crude locomotion, suggesting that further developments of these neurons are required if they are to contribute to the development of locomotor behaviors.

Hindbrain V2a neurons with spinal projections are more likely to play roles in the development of the locomotor repertoire, therefore we examined the times of birth of these neurons in further detail. First we focused on large spinal projecting neurons (reticulospinal neurons) that were labeled by the dye injection in the spinal cord (Figure 2C-D, n = 4 for each timepoint). These neurons are identifiable across animals and are named based on their anatomical features (Kimmel, Powell, and Metcalfe 1982; Mendelson 1986). The dorsal subpopulation of reticulospinal neurons (MiD2i, MiD3i, RoM2, RoM3, dorsal MiV1 (see Methods)) corresponded to V2a neurons born by 24 hpf (Figure 2C), while the majority of ventral reticulospinal neurons (RoV3 and ventral MiV1) corresponded to V2a neurons born by 36 hpf (Figure 2D). This birth order is consistent with previous birthdating analysis of reticulospinal neurons based on the incorporation of a marker into replicating DNA (Mendelson 1986). Other than the large spinal projection neurons that are labeled consistently by dye injection, a majority of the V2a neurons in the caudal hindbrain have also been shown to project to the spinal cord (Kimura et al. 2013). In this region, the cell bodies of the earliest-born V2a neurons were located lateral to the populations born afterwards (Figure 2 E). This clear topographical organization allows us to readily identify the earliest-born population and the later-born population in the caudal hindbrain. Notably, this region contained the youngest group of neurons among the hindbrain V2a descending neurons we examined, suggesting their roles in the refined locomotion that develops later (Figure 2E, 48 hpf conversion).

**Figure 2.**
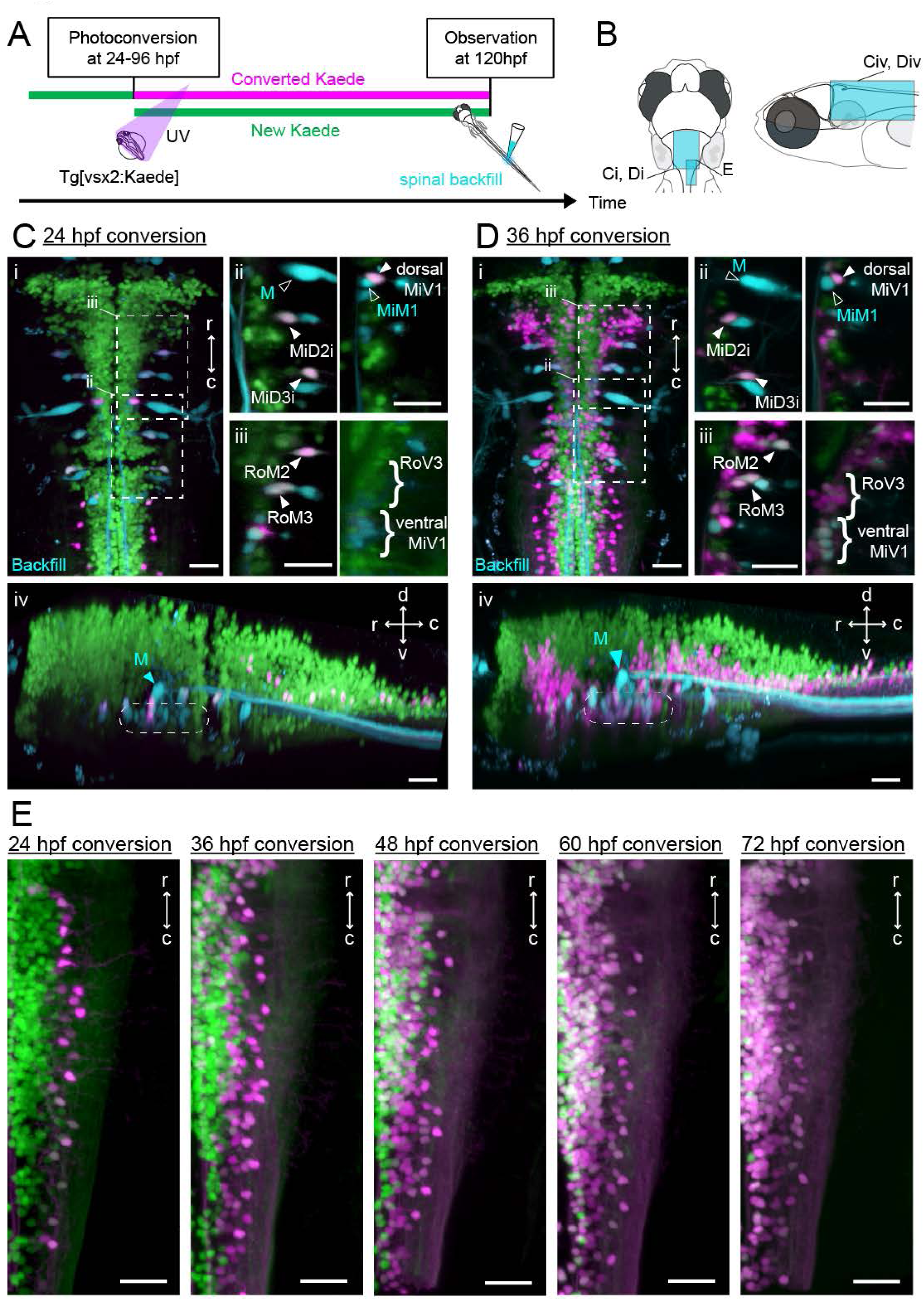
Birthdating and identification of hindbrain V2a descending neurons. **A.** Experimental procedure. **B.** Regions displayed (cyan patch). **C.** Hindbrain V2a neurons photoconverted at 24 hpf with spinal backfill of reticulospinal neurons. Green, unconverted Kaede; magenta, photoconverted Kaede; cyan, backfill; M, Mauthner. (i) Top-down view. Dotted rectangles indicate the locations of panels ii and iii. (ii) Optical slices showing caudal reticulospinal neurons. (iii) Optical slices showing rostral reticulospinal neurons (left panel, dorsal slice; right panel, ventral slice). Filled arrowheads and curly brackets indicate V2a reticulospinal neurons. Open arrowheads indicate non-V2a reticulospinal neurons. (iv) Side-view. Dotted rounded rectangle indicates the location of ventral reticulospinal neurons. **D.** Hindbrain V2a neurons photoconverted at 36 hpf, and reticulospinal neurons labeled with spinal backfill. Panels are organized as in C. (i) Top-down view. (ii) Optical slices showing caudal reticulospinal neurons. (iii) Optical slices showing rostral reticulospinal neurons (left panel, dorsal slice; right panel, ventral slice). (iv) Side-view. Dotted rounded rectangle indicates the location of ventral reticulospinal neurons. **E.** Top-down views of caudal hindbrain V2a neurons photoconverted at 24, 36, 48, 60, and 72 hpf. r, rostral; c, caudal; d, dorsal; v, ventral. Scale bars, 30 um.

For the hindbrain V2a neurons to contribute to the development of the locomotor repertoire, they first need to establish connections to spinal neurons. To examine the development of spinal projections we optically back labeled V2a descending neurons from the spinal cord (‘optical backfill’) using photoconversion restricted to the rostral spinal cord (Kimura et al. 2013). This procedure was repeated at multiple developmental time points to examine the developmental sequence of spinal projections (Figure 3A, n=6 for each timepoint). The optical backfill at 36 hpf labeled the dorsal V2a but not the ventral V2a reticulospinal neurons (Figure 3C) while the backfill at 60 hpf labeled both the dorsal and ventral reticulospinal neurons (Figure 3D), indicating that these neurons had projected to the rostral spinal cord in sequence based on their respective differentiation times. The V2a subpopulation in the caudal hindbrain showed a pattern similar to that indicated by the birthdating analysis (Figure 3E). This indicates that the sequential development of spinal projections based on differentiation time holds true for the caudal hindbrain V2a subpopulation as well. Among these descending neurons, the caudal medial V2a neurons were the last to project to the spinal cord. As these neurons had descending processes in the rostral spinal cord by 84 hpf, it is plausible that they contribute to the refined locomotion that appears as early as 96 hpf.

**Figure 3.**
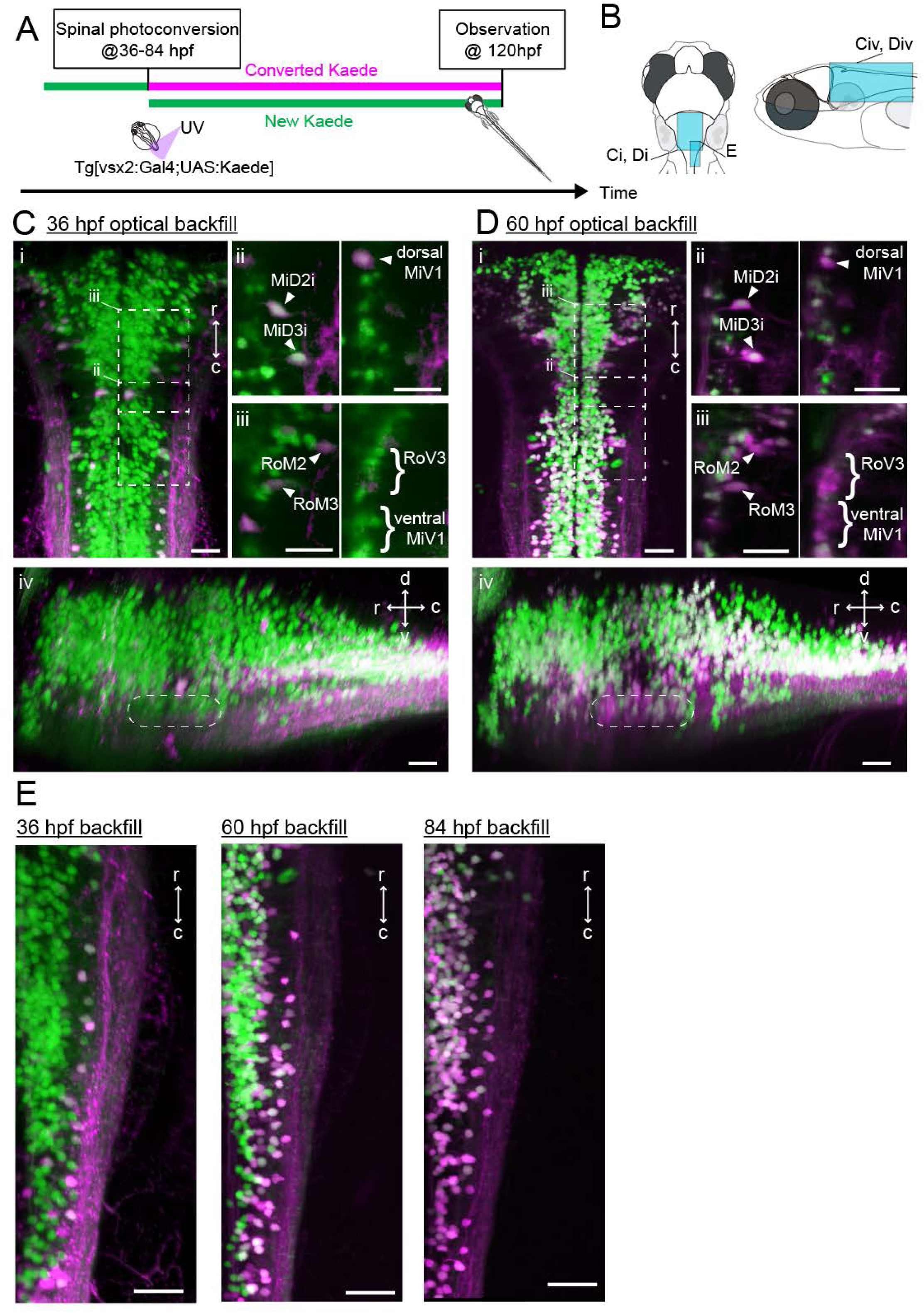
Development of spinal projections of hindbrain V2a descending neurons. **A.** Experimental procedure. **B.** Regions displayed. **C.** Hindbrain V2a neurons optically backfilled from the rostral spinal cord at 36 hpf. Magenta, photoconverted Kaede; green, unconverted Kaede. (i) Top-down view. Dotted rectangles indicate the locations of panels ii and iii. (ii) Optical slices showing caudal V2a reticulospinal neurons. (iii) Optical slices showing rostral V2a reticulospinal neurons (left panel, dorsal slice; right panel, ventral slice). (iv) Side-view. Dotted rounded rectangle indicates the position of ventral reticulospinal neurons. **D.** Hindbrain V2a neurons optically backfilled from rostral spinal cord at 60 hpf. Photoconverted Kaede is shown in magenta and unconverted Kaede in green. (i) Top-down view. Dotted rectangles indicate the locations of panels ii and iii. (ii) Optical slices showing caudal reticulospinal neurons. (iii) Optical slices showing rostral reticulospinal neurons (left panel, dorsal slice; right panel, ventral slice). (iv) Side-view. Dotted rounded rectangle indicates the location of ventral reticulospinal neurons. **E.** Top-down views of caudal hindbrain V2a neurons optically backfilled at 36, 60 and 84 hpf. r, rostral; c, caudal; d, dorsal; v, ventral; Scale bars, 30 um.

Collectively, the results of our developmental analysis showed the timeline of neurogenesis and spinal projections of hindbrain V2a descending neurons relative to the development of the locomotor repertoire. Furthermore, we found that the birth order of these neurons dictated the positions of their cell bodies and the order of their spinal projections. Most importantly, this analysis established a way to identify the birthdates of these neurons based on the positions of their cell bodies. This set the stage for functional analysis of these neurons, wherein we characterized how neurons of different age were recruited during the distinct locomotor patterns that fish develop in sequence so as to examine the functional links between the development of V2a neurons and the development of locomotor behaviors.

### 2. Hindbrain V2a neurons of different ages are recruited differentially during distinct locomotor patterns

To identify functional cell groups related to the crude and refined locomotor behaviors that appear in sequence during development, we examined the recruitment of hindbrain V2a neurons expressing the calcium indicator GCaMP6s (Chen et al. 2013) using whole-hindbrain two-photon volumetric imaging at 2 Hz in 120-132 hpf fish, which are capable of generating both forms of locomotion. Fish were paralyzed and axial motor activity was monitored using glass pipettes attached to motor nerves innervating axial muscles on both sides (Figure 4; Supplementary video 1). We used two stimulus paradigms to produce distinct tactile-induced crude fictive locomotion patterns (6 fish for each paradigm). The first paradigm made use of a transient electrical pulse delivered to one side of the head, which reliably led to axial motor activity with rhythmic bursting that was distinctively stronger and faster than the rhythmic bursting in spontaneously occurring axial motor activity (Figure 4A i). These episodes started with axial motor activity on the side contralateral to the stimulation, indicating that they were escapes away from the stimulus (Figure 4A i). The second paradigm employed a gradual mechanical stimulus applied to the front of the head, which typically led to axial motor activity with strong but slow rhythmic bursting that was (Figure 4B i) similar to the “struggling” behavior reported previously (Liao and Fetcho 2008). In both paradigms, spontaneous axial motor activity exhibited weak and slow rhythmic bursting, suggesting that these episodes corresponded to the more refined locomotor pattern observed spontaneously in unparalyzed condition. Peripheral motor nerve recordings from a single site on each side did not allow us to determine if the rostrocaudal patterns of axial motor activity were organized similarly to those observed in unparalyzed fish, but the amplitude and left-right alternation frequency of bursting axial motor activity were similar to those observed in unparalyzed fish.

**Figure 4.**
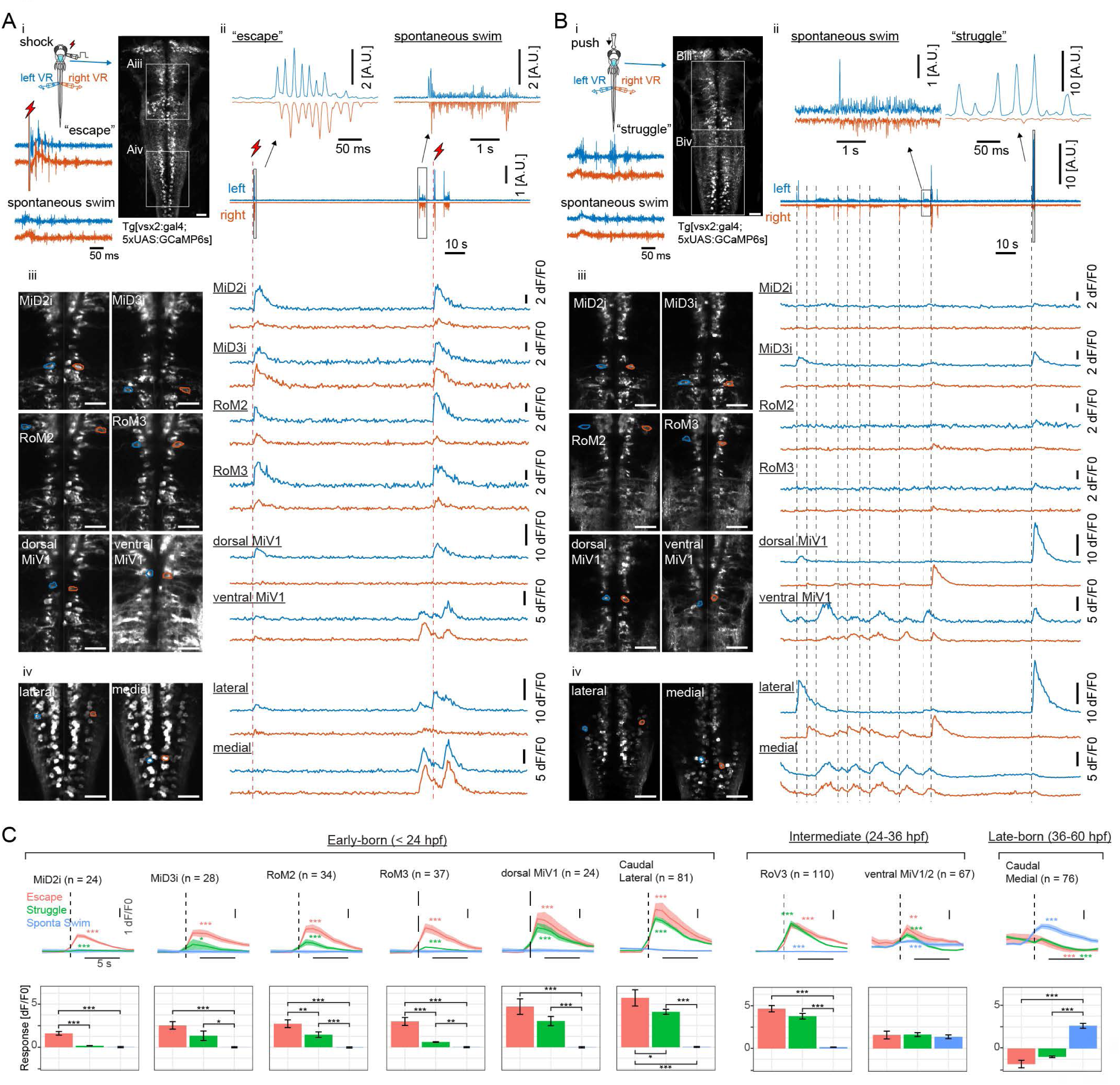
Recruitment of hindbrain V2a neurons during distinct locomotor behaviors. **A.** Ca^2+^ responses of hindbrain V2a neurons during putative escape induced by a transient electrical stimulus to the head. (i) Experimental setup. The image on the right shows the top-down view of hindbrain V2a neurons expressing GCaMP6s. White squares indicate the regions displayed in subsequent panels. Example raw traces from left and right ventral root (VR) activity during a putative escape and spontaneous swim episode (blue trace, left VR signal; orange trace, right VR signal; red thunder symbol, onset of an electrical pulse). (ii) Example traces of processed swim signals during a spontaneous weak swim episode and shock-induced strong swim episode, (blue trace, processed left VR; orange trace, processed right VR; red thunder mark, shock onset). (ii-iii) Location of hindbrain V2a neurons and corresponding Ca^2+^ traces during the locomotion patterns shown in panel i (blue, left ROIs and their Ca^2+^ responses; orange, right ROIs and their Ca^2+^ responses). Vertical dotted red lines indicate the onsets of the shock stimuli. (ii) Ca^2+^ traces of rostral V2a neurons (iii) Ca^2+^ traces of caudal hindbrain V2a neurons. (Scale bars in the images, 30 um). **B.** Ca^2+^ responses of hindbrain V2a neurons during a putative struggle induced by the gradual mechanical stimulus. The panels are organized as in A. Note the strong and slowly alternating (between left and right) axial motor activity in panel i. **C.** Locomotion event-triggered average Ca^2+^ responses of hindbrain V2a neurons sorted based on their birthdate. Top, Time course of the event-triggered average Ca^2+^response (Orange trace, putative escape; Green trace, putative struggle; Blue trace, spontaneous slow swim). Shaded area indicates standard error across replicates for each cell type (n, number of cells). Significant changes from the baselines are indicated with asterisks (paired t-test, \**P* < 0.05, \*\**P* < 0.01, \*\*\**P* <0.001). Bottom, Peak amplitude of locomotion event-triggered average Ca^2+^ response (Orange bar, putative escape; Green bar, putative struggle; Blue bar, spontaneous slow swim). Significant differences in response amplitude across conditions are indicated with asterisks (Tukey test, \**P* < 0.05, \*\**P*<0.01, \*\*\**P*< 0.001).

To examine neuronal recruitment, we first analyzed imaging data during the two stimulus paradigms (Figure 4A, B) focusing on the V2a descending neurons we had identified based on development. In the electrical shock paradigm (Figure 4A), almost all of the identifiable dorsal neurons were recruited during putative escapes but primarily on the side ipsilateral to the leading axial motor activity (Figure 4A iii, blue traces). This is consistent with the primarily ipsilateral projections of these neurons (Kinkhabwala et al. 2011; Kimmel, Powell, and Metcalfe 1982). On the other hand, the ventral MiV1 and caudal medial neurons showed stronger activity during weak spontaneous swimming. In the gradual mechanical stimulus paradigm, a subset of the neurons recruited during putative escapes were recruited when there was strong “struggle” like motor activity on the ipsilateral side (Figure 4B iii-iv, blue traces close to the end). The ventral MiV1 and caudal medial cells were again recruited during weak spontaneous swimming. When we sorted these cell types based on their birthdate and examined their responses during these motor patterns, it became apparent that there was a systematic relationship between recruitment and birthdate (Figure 4C, 6 fish for each paradigm). The early-born group (Figure 4C, <24 hpf) did not show any clear response during spontaneous weak swimming, but many neurons in this group showed significant responses during putative struggles induced by the gradual mechanical stimulus. The responses in a few of these neurons were even stronger during putative escapes elicited by the transient electrical pulse stimulus (Figure 4C, MiD2i, RoM2 and RoM3). On the other hand, the intermediate-aged group (Figure 4C, 24-36 hpf) showed mixed recruitment patterns; the subgroup in the rostral hindbrain (Figure 4C, RoV3) showed comparably strong activity during putative escapes and putative struggles but not during weak spontaneous swims whereas the subgroup in the middle hindbrain (Figure 4C, ventral MiV1/2) showed comparably strong activity in all three conditions. In contrast, the late-born group in the caudal hindbrain (Figure 4C, 36-60 hpf) showed more activity only during weak spontaneous swimming. Interestingly, during stimulus-evoked strong swims, these neurons showed decreased activity instead, raising the possibility of inhibition during strong swims. Taken together, our findings indicate that a recruitment pattern that is based on the time of differentiation holds true for all the hindbrain V2a descending neurons that we identified based on development; the early-born group is recruited during the crude forms of locomotion that appear early in development whereas the late-born group is recruited during the refined locomotion that appears later in development. This finding, along with the finding that spinal projections develop sequentially, suggests that distinct functional groups emerge in developmental sequence and contribute to the development of locomotor behaviors. Furthermore, the active set of hindbrain V2a neurons switched based on the type of ongoing motor activity, which is reminiscent of the recruitment of spinal interneurons (McLean et al. 2008). This raises the possibility that distinct functional groups of hindbrain V2a neurons may provide excitatory drive to corresponding spinal functional groups.

To see if our observations about neuronal recruitment extend to the complete set of hindbrain V2a neurons, we performed regression analysis (Miri, Daie, Burdine, et al. 2011) (Figure 5). In the electrical pulse paradigm, we used two regressors to distinguish the activity related to shock-induced putative escapes from activity related to weak spontaneous swimming (Figure 5A ii). The activity map for the shock-evoked putative escapes (Figure 5A iii, left; Figure 5A iv, magenta) revealed a large population of recruited neurons on the side contralateral to the stimulus (ipsilateral to the leading side of motor activity), consistent with their primarily ipsilateral projections (Kinkhabwala et al. 2011). This includes the dorsal early-born group and the rostral intermediate group (RoV3) in the rostral hindbrain (Figure 5A V, magenta) and a large number of the lateral early-born neurons in the caudal hindbrain (Figure 5A vi, magenta). The activity map for spontaneous weak swimming revealed the middle intermediate group (ventral MiV1/2) in the rostral hindbrain (Figure 5 A iii, right) and a large number of the medial late-born neurons in the caudal hindbrain (Figure 5A iii, right; Figure 5A vi, cyan). For the gradual mechanical stimulus paradigm that produced putative struggles that varied in latency, we constructed three regressors based on the strength of the axial motor activity: two regressors for the strong motor activity for each side to capture the struggle related activity and one regressor for the weak motor activity to capture the activity related to spontaneous weak swims (Figure 5B ii). The activity maps for the strong motor activity (Figure 5B iii, left and right; Figure 5B iv, yellow and magenta) revealed a large population of neurons ipsilateral to the side of motor activity (Figure 5B iii, left and right) consisting of a subpopulation of the dorsal early-born group, the rostral intermediate group (RoV3) in the rostral hindbrain (Figure 5B v), and a large number of the early-born lateral neurons in the caudal hindbrain (Figure 5B vi, magenta). The map for spontaneous swimming revealed the same set of the neurons shown for spontaneous swimming occurring in the electrical pulse paradigm (Figure 5B iii, middle; Figure 5B iv, cyan). These regression maps were reproducible across fish (Figure 5 supplement 1) and further support the differentiation time dependent recruitment pattern revealed by the ROI analysis. Note that the additional recruitment of rostral V2a neurons during shock-induced fast and strong swims became more obvious with this analysis (Fig 5 supplement 1B). Interestingly, these maps revealed a striking functional separation of neuropil in the caudal hindbrain (Figure 5A vi; Figure 5B vi; Figure 5 supplement 1C). The neuropil active during weak spontaneous swimming was located lateral to the neuropil active during putative struggles (Figure 5B vi, filled arrowheads; Figure 5 supplement 1C, Struggle, arrowheads). This functional segregation is the opposite of the functional segregation of cell bodies in the caudal hindbrain (Figure 5B vi, open arrowheads). This neuropil separation was maintained when additional cells in the rostral hindbrain were recruited for the shock-induced strong and fast swim (Figure 5A vi, filled arrowheads; Figure 5 supplement 1C, Escape, arrowheads), suggesting that these additional cells also followed this functional separation. Over all, this raises the possibility that these functional neuronal groups project their descending axons through distinct regions in the neuropil.

**Figure 5.**
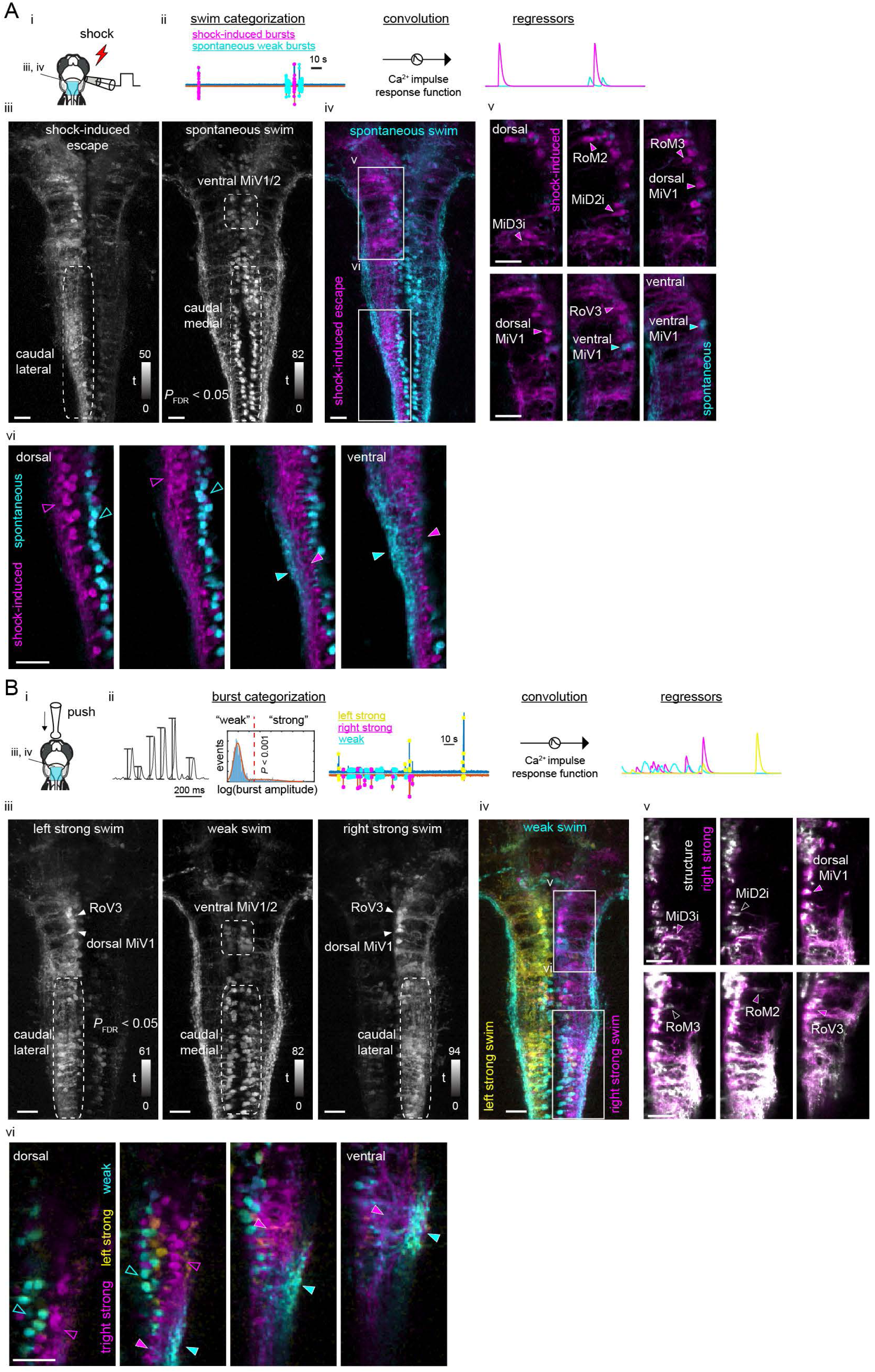
Functional segregation of the neuropil of hindbrain V2a neurons as revealed by regression analysis. **A.** Regression analysis of hindbrain V2a neurons in the transient electrical stimulus (shock) experiment. (i) Experimental setup and the region displayed (cyan patch). (ii) Regressors used to create activity maps (see Methods). (iii) Top-down view of activity maps for each regressor. T-value was coded in grayscale and thresholded at *P_FDR(False Discovery Rate_* < 0.05 (see Methods). Dotted round rectangles indicate distinct caudal hindbrain populations revealed by each regressor. (iv) Overlay of activity maps (cyan, spontaneous swim; magenta, shock-induced fast swim). Rectangles indicate the locations of the images in panel v and vi. (v) Optical slices of the overlaid activity maps in the rostral hindbrain (magenta, shock-induced swim; cyan, spontaneous swim). Arrowheads indicate reticulospinal neurons identified in each map. (vi) Optical slices of the overlaid activity maps in the caudal hindbrain (magenta, shock-induced swim; cyan, spontaneous swim). Arrowheads highlight the segregation of spontaneous swim related signal (cyan open arrowhead, cell bodies; cyan filled arrowhead, neuropil) and shock-induced swim related signal (magenta open arrowhead, cell bodies; magenta filled arrowhead, neuropil). Scale bars, 30 um. **B.** Regression analysis of hindbrain V2a neurons in the gradual mechanical stimulus (push) experiment. (i) Experimental setup and the region displayed (cyan patch). (ii) Regressors used to create activity maps (see Methods). (iii) Top-down view of activity maps for each regressor. T-value was coded in grayscale and thresholded at *P_FDR_* < 0.05. Dotted round rectangles and arrowheads are used to highlight structures revealed in each activity map. (iv) Overlay of activity maps (cyan, weak swim; yellow, left strong swim; magenta, right strong swim). Rectangles indicate the locations of the images in panel v and vi. (v) Optical slices of the activity map for the right strong swim (magenta) overlaid on the structural image of V2a neurons (white) in the rostral hindbrain (magenta arrowheads, recruited reticulospinal neurons; white open arrowheads, non-recruited reticulospinal neurons). (vi) Optical slices of the overlaid activity maps in the caudal hindbrain (magenta, right strong swim; cyan, weak swim). Arrowheads highlight the segregation of weak swim related signal (cyan open arrowhead, cell bodies; cyan filled arrowhead, neuropil) and strong swim related signal (magenta open arrowhead, cell bodies; magenta filled arrowhead, neuropil). Scale bars, 30 um.

To sum up, we showed a correspondence between the birth order of hindbrain V2a neurons and their involvement in distinct forms of locomotion that appear in sequence. This suggests that distinct functional cell groups emerge in the hindbrain and project to the spinal cord in sequence and contribute to the diversification of locomotor repertoire. Furthermore, the similarity in the recruitment of hindbrain and spinal functional groups as well as the spatial segregation of the neuropil of the hindbrain functional groups both raise the possibility that parallel pathways connect matching sets of hindbrain and spinal functional groups. Having established a link between the birth order and function of hindbrain V2a neurons, we next examined the spinal organization of descending processes from hindbrain functional groups.

### 3. Chronological layering of spinal projections of hindbrain V2a neurons suggests parallel descending pathways organized by birthdate

To understand how the spinal projections from hindbrain V2a neurons are organized to support the development of the locomotor repertoire, we examined if the aforementioned sequentially generated functional groups of hindbrain V2a neurons exhibit distinct innervation patterns to the spinal cord. First, we examined the overall neuropil organization of hindbrain and spinal V2a neurons based on the time of differentiation by photoconverting early-born V2a neurons at 24 and 36 hpf (Figure 6A, n = 6 fish for each timepoint). In the hindbrain (Figure 6A iii), and throughout the spinal cord (Figure 6A iv-vi) the neuropil from V2a neurons born before 24 hpf (Figure 6A iii-vi, magenta arrowheads) was located medial to the neuropil from neurons born after 24 hpf (Figure 6A iii-vi, green arrowheads), consistent with the functional segregation of neuropil we observed in the caudal hindbrain. The neuropil from V2a neurons born before 36 hpf was still located medial to the neuropil from neurons born after 36 hpf (Figure 6A viii-xi, arrowheads). However, it extended more towards the lateral surface than the neuropil from V2a neurons born before 24 hpf (Figure 6A xii). This suggested that axonal processes from new neurons were continuously being added laterally to preexisting processes, forming layers based on the time of differentiation.

**Figure 6.**
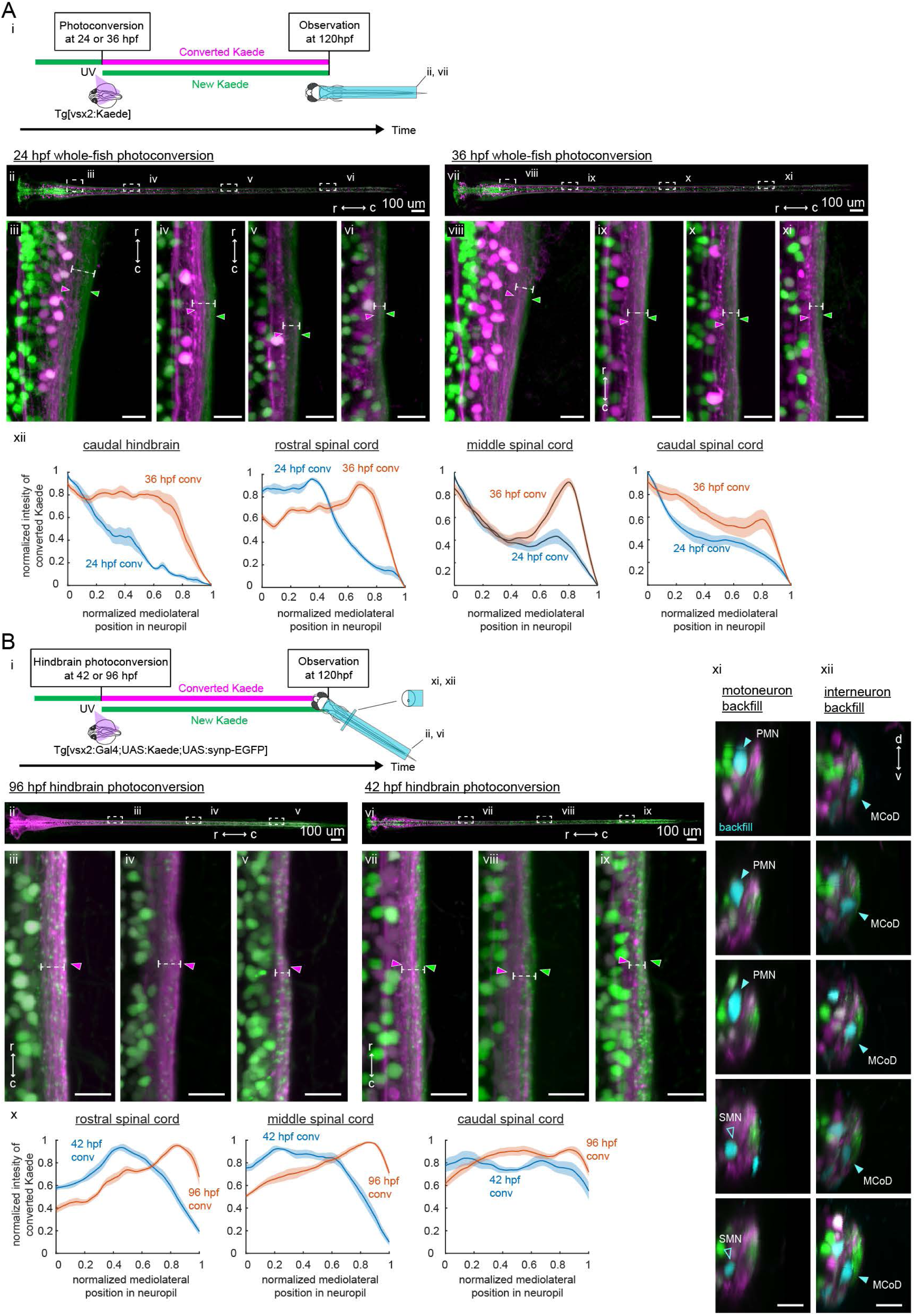
Birthdate-related segregation of spinal projections from hindbrain V2a neurons. **A.** Birthdate-related segregation of the neuropil of V2a neurons in the hindbrain and the spinal cord. (i) Experimental procedure and region displayed (gray patch). Converted Kaede is shown in magenta and unconverted Kaede is shown in green. (ii-vi) Segregation of neuropil from V2a neurons born before and after 24 hpf. (ii) Top-down view of the hindbrain and the spinal cord. Dotted rectangles indicate the locations of the images in the following panels. (iii-vi) Close up views of the early-born neuropil (magenta arrowheads) and the late-born neuropil (green arrowheads) in the hindbrain and spinal cord. Dotted lines indicate the mediolateral extent of neuropil. (iii) Top-down view of the caudal hindbrain. (iv) Top-down view of the rostral spinal cord. (v) Top-down view of the middle spinal cord. (vi) Top-down view of the caudal spinal cord. r, rostral; c, caudal. Scale bars, 15 um. (vii-xi) Neuropil segregation between V2a neurons born before and after 36 hpf. Panels are organized as in ii-vi. (xii) Intensity profile of photoconverted Kaede in the neuropil. Normalized intensity of photoconverted Kaede is plotted as a function of normalized mediolateral position (dashed white lines in iii-vi, viii-xi) in the neuropil. Lines indicate the average of intensity distributions across fish, while shaded areas around the line represent the standard error (n = 6). Blue, photoconversion at 24 hpf; orange, photoconversion at 36 hpf. **B.** Segregation of descending projections from hindbrain V2a neurons. (i) Procedure of the experiments and location of the images in panel ii and vi. (magenta, photoconverted Kaede; green, unconverted Kaede). (ii-v) Hindbrain V2a neurons are photoconverted at 96 hpf to highlight the spinal projections of all hindbrain V2a descending neurons. Magenta arrowheads indicate the spinal neuropil filled with the hindbrain V2a descending neurons. Dotted lines indicate the mediolateral extent of neuropil. (ii) Top-down view of hindbrain and spinal cord. Dotted rectangles indicate the location of the images in panel iii-v. (iii) Top-down view of the rostral spinal cord, (iv) Top-down view of the middle spinal cord. (v) Top-down view of the caudal spinal cord. (vi-xii) Hindbrain V2a neurons are photoconverted at 42 hpf to highlight the spinal projections of the early-born and intermediate-aged hindbrain V2a neurons. Magenta arrowheads indicate the neuropil region occupied by the early-born hindbrain V2a neurons. Green arrows indicate spinal neuropil regions that have fewer projections of the early-born hindbrain V2a neurons. Dotted lines indicate the mediolateral extent of neuropil. Panels vi to ix are organized as in panels ii-v. (x) Intensity profile of photoconverted Kaede in the neuropil. Normalized intensity of photoconverted Kaede is plotted against normalized mediolateral position in the neuropil (dashed white lines in iii-v, vii-ix). Lines indicate the average of intensity distributions across fish, while shaded areas around the line represent the standard error (n = 8). Blue, hindbrain photoconversion at 42 hpf; orange, hindbrain photoconversion at 96 hpf. (xi) Coronal views of spinal motoneurons in the rostral spinal cord. One side of the spinal cord is shown. Backfilled motoneurons are shown in cyan. Closed cyan arrowheads indicate primary motoneurons (PMN). Open arrowheads indicate secondary motoneurons (SMN). (xii) Coronal views of spinal interneurons in the rostral spinal cord. One side of the spinal cord is shown. Backfilled Interneurons are shown in cyan. Closed cyan arrowheads indicate the late-born spinal interneuron, multipolar commissural neuron (MCoD).r, rostral; c, caudal; d, dorsal; v, ventral. Scale bars, 15 um.

To see if this neuropil organization was maintained in the descending processes of hindbrain V2a neurons, we photoconverted V2a neurons only in the hindbrain (Figure 6B). The Gal4-UAS system was used to gain enough expression of Kaede to be able to visualize the spinal processes of hindbrain V2a neurons. Putative presynaptic terminals were also labeled with synaptophysin-GFP driven by the Gal4-UAS system. To visualize the spinal projections from almost all the age groups of the hindbrain V2a neurons, we photoconverted Kaede in the hindbrain at 96 hpf and imaged them at 120 hpf (Figure 6B i, n = 8 fish). The descending processes from hindbrain V2a neurons covered almost all of the neuropil region formed by V2a neurons (Figure 6B iii-v, magenta arrowheads). To visualize the spinal projections from only the early-born and intermediate-aged hindbrain V2a groups, UV light was shone on the hindbrain at 42 hpf (n = 8 fish). This conversion time was selected to take account of the delay in the expression of Kaede driven by the Gal4-UAS system and to match the age groups labeled by the photoconversion of Tg(vsx2:Kaede) at 36 hpf. The descending processes from the early-born and intermediate-aged groups also reached the caudal part of the spinal cord (Figure 6B ix) and were in the medial part of the neuropil in the rostral and middle spinal cord (Figure 6B vii-viii, magenta arrowheads). Comparison of the lateral extent of the descending processes between the early-born and intermediate-aged groups (Figure 6B x, blue lines) and all groups (Figure 6B x, orange lines) indicated that the descending processes from the late-born group were most prominent in the lateral neuropil in the rostral and middle spinal cord. Taken together, these results indicated that the spinal projections from hindbrain V2a neurons were also layered based on the time of differentiation and that the late-born group had shorter descending processes than the early-born and intermediate-aged groups. The difference in the axon length across different age groups is consistent with the pattern previously revealed by single-cell labeling (Kinkhabwala et al. 2011). These observations support the notion that the sequentially generated hindbrain functional groups provide parallel spinal pathways.

To examine the pattern of connections made by each pathway onto possible postsynaptic spinal neurons, we highlighted the descending processes of early-born and intermediate-aged hindbrain V2a neurons by photoconversion in the hindbrain, and additionally backfilled the following two spinal cell types with a far-red dye. The first cell type is primary motoneurons (PMN) which are among the earliest-born neurons in the spinal cord (Myers, Eisen, and Westerfield 1986) (Figure 6B xi, n = 8 fish). They are only recruited during the strongest of movements in contrast to secondary motoneurons (SMN) which are also recruited during weaker movements (McLean et al. 2007; Menelaou and McLean 2012; Wang and Brehm 2017). The second cell type is multipolar commissural neurons (MCoD), which are later-born excitatory Interneurons in the rostral spinal cord (Figure 6B xii, n = 8 fish). They are only active during slow swimming and have monosynaptic connections to SMNs in the caudal spinal cord (McLean et al. 2008; Fetcho and McLean 2010). We found that PMNs were located close to the medial neuropil region occupied by the spinal projections from the early-born hindbrain V2a population, whereas MCoDs were located close to the lateral neuropil region occupied by the spinal projections from the late-born population. Thus, these results support the notion that there are parallel pathways between the hindbrain and the spinal cord that are separable by differentiation time.

The innervation patterns exhibited by these hindbrain V2a age groups are consistent with the type of axial muscle activity observed during the locomotor patterns that these age groups participate in: direct activation of PMNs throughout the spinal cord by the early-born group is expected to lead to the fast and powerful whole-body bends observed during the crude and strong forms of locomotion whereas indirect activation of SMNs in the caudal spinal cord through MCoDs should lead to the slow bend of the caudal tail observed during the more refined and weaker locomotion. Thus, these observations raise the possibility that each age group contributes to the kinematics specific to the the locomotor pattern in which this group participates.

### 4. In-depth analyses of V2a reticulospinal neurons

So far we experimentally investigated the overall organization of the hindbrain V2a population and the results suggest that birthdate-dependent parallel circuit organization underlies the development of locomotor behaviors. To examine this more rigorously and to gain deeper insights about the biophysical properties of V2a neurons and their descending pathways, we examined various subsets of V2a reticulospinal neurons in depth.

#### 4-1. Neuronal excitability and synaptic inputs in V2a pathways vary in accordance with birthdate-related recruitment pattern

To electrophysiologically examine the birthdate-related recruitment pattern, we focused on MiV1 neurons, a small cluster of V2a reticulospinal neurons in rhombomere 4 (Figure 7A), which included neurons varying in birthdate and recruitment pattern (Figure 2, 4, 5).

**Figure 7.**
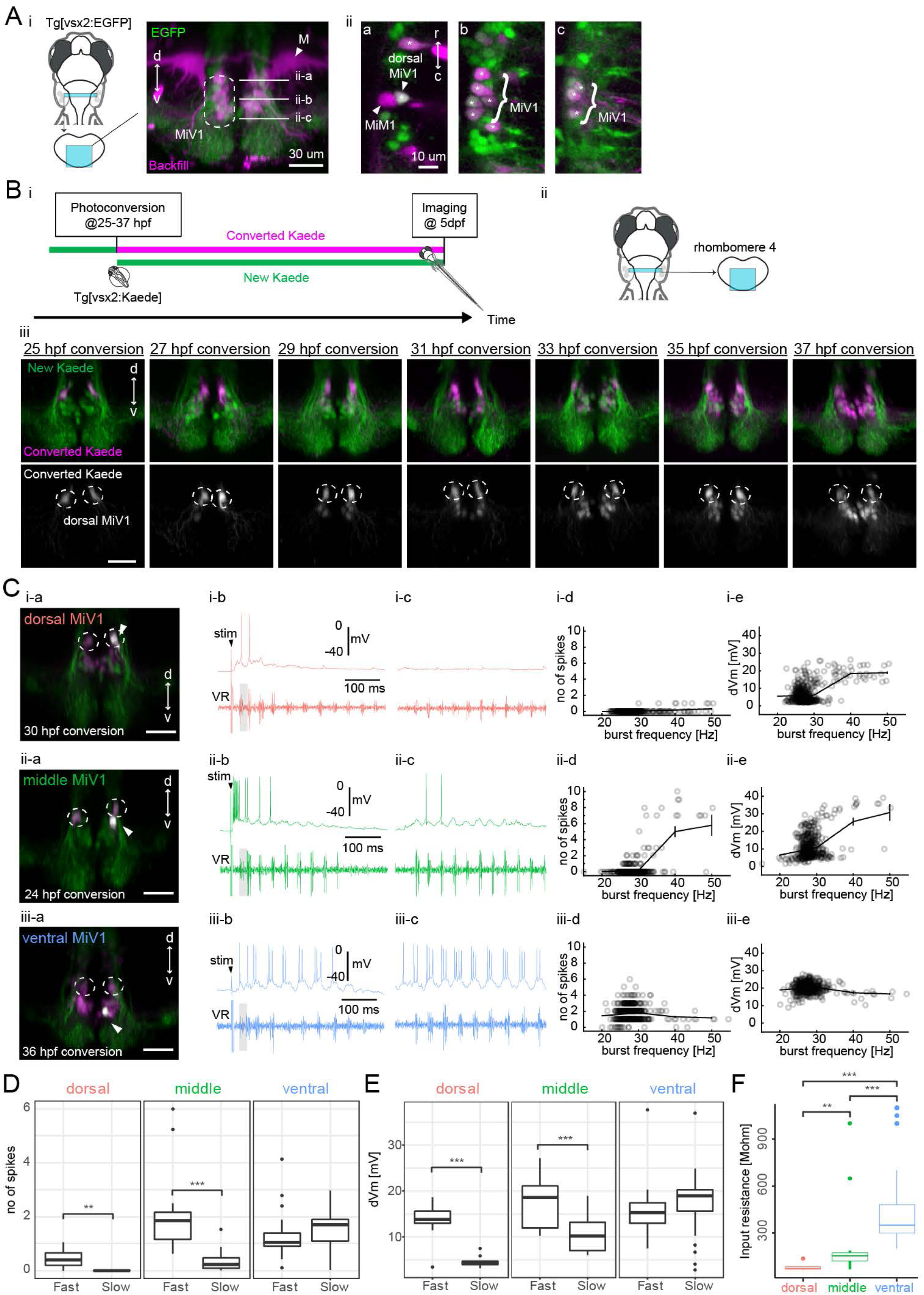
Birthdating and electrophysiological analysis of V2a reticulospinal neurons in rhombomere 4. **A.** V2a reticulospinal neurons in rhombomere 4. (i) Coronal view of V2a reticulospinal neurons in the region of rhombomere 4 shown by the cyan patch (green, EGFP; magenta, backfill; d, dorsal; V, ventral). Mauthner cell (M) and MiV1 neurons are highlighted. White horizontal lines in the image indicate the optical slices shown in panel ii. (ii) Optical slices showing reticulospinal neurons at different depths indicated by white lines in i. Asterisks indicate backfilled V2a neurons. MiM1 and dorsal MiV1 are highlighted with arrowheads. Other more ventral MiV1s are highlighted with curly brackets. **B.** Birthdate-related topographical organization of V2a RS neurons in rhombomere 4. (i) Experimental procedure (ii) Region displayed (cyan patch) (iii) Coronal views of V2a RS neurons in rhombomere 4 at 5 dpf showing Kaede photoconverted at a specific time point (25-37 hpf) in magenta in upper panels and gray in lower panels. Dotted circles indicate dorsal MiV1. **C.** Whole-cell recordings of V2a reticulospinal neurons in rhombomere 4. (i) Recruitment of an early-born V2a neuron (dorsal MiV1). (i-a) Coronal view of the patched cell (gray). Green, unconverted Kaede. Magenta, converted Kaede. (i-b) Intracellular activity during shock-induced fast swimming. An arrowhead indicates the onset of the tail shock. VR, ventral root recording. Gray shaded boxes indicate similar fast burst frequencies. (i-c) Intracellular activity during spontaneous slow swimming. (i-d) Number of spikes per cycle as a function of burst frequency. Open circles are raw data points. The solid line represents mean ± standard error from data binned at 10-Hz intervals. (i-e) Membrane depolarization as a function of burst frequency. (ii) Data from an intermediate V2a neuron (middle MiV1) are organized as in i. (iii) Data from a late-born V2a neuron (ventral MiV1) are organized as in i. **D.** Number of spikes per cycle during fast burst frequency (>35 Hz) and slow burst frequency (<35 Hz) for each age group (dorsal MiV1, n = 16; middle MiV1, n = 26; ventral MiV1, n = 46). Significant differences are indicated with asterisks (*** *P* < 0.01, \*\*\**P* < 0.001). **E.** Membrane depolarization during fast swim (>35 Hz) and slow burst frequency (<35 Hz) for each age group (dorsal MiV1, n = 16; middle MiV1, n = 26; ventral MiV1, n = 46). Significant modulations based on burst frequency are indicated with asterisks (\*\*\**P* < 0.001). **F.** Input resistance for each age group (dorsal MiV1, n = 16; middle MiV1, n = 26; ventral MiV1, n = 46). Significant differences are indicated with asterisks (Dunn”s test, \*\**P* < 0.01, *** *P* < 0.001).

First, we examined the spatial organization of neurons in this cluster based on the time of differentiation, as indicated by photoconversion-based birthdating from 25 hpf to 37 hpf (Figure 7B, n = 6 fish for each timepoint). We found that newly generated V2a neurons were systematically located ventral to preexisting ones (Figure 7B iii), in contrast to a previous study reporting that late-born neurons are located dorsal to early-born neurons except in rhombomere 6 (Kinkhabwala et al. 2011). To see if this inverted organization was due to migration, we performed time lapse imaging from 25 hpf to 37 hpf (Figure 7 supplement 1A, n = 6) and from 48 hpf to 78 hpf (Figure 7 supplement 1B, n = 6). We found that newly generated neurons were indeed initially located dorsal to preexisting ones (Figure 7 supplement 1A iii) but the subpopulation of neurons born before 48 hpf migrated ventrally after 48 hpf (Figure 7 supplement 1B iii, magenta cells; Supplementary video 2). This supported the idea that the inverted organization we observed in this region was indeed due to radial migration. This analysis revealed a fine-scale birthdate-related topographical organization, which our previous analyses overlooked, and provided an opportunity to examine in detail the relationship between ontogeny and recruitment pattern.

To examine the activity of MiV1 neurons during locomotion, we performed whole-cell recordings in a paralyzed fictive swim preparation (Figure 7C-F). We delivered electrical shocks to the tail to induce putative escapes that showed fast frequency of bursting activity in axial motor recording (Figure 7C b). Locomotor episodes with slower frequency of bursting activity were observed spontaneously (Figure 7C c) as in the functional imaging experiments. However, whole-cell recordings allowed us to examine how activity changes in relationship to burst frequency within each swim episode at the level of spiking and subthreshold activity. The dorsally located early-born MiV1 neurons (n = 16) showed strong depolarization leading to action potentials during the fastest phase of fast swim episodes evoked by electrical shock to the tail (Figure 7C i-b, shaded cycle). However, they showed no clear depolarization during spontaneous slow swim episodes (Figure 7C i-c). The middle MiV1 neurons that belong to the intermediate age group (n = 26) showed clear spiking activity at the fastest phase of the shock-induced fast swimming. They also showed clear rhythmic depolarization leading to occasional firing during spontaneous slow swimming (Figure 7C ii). The ventral late-born MiV1 neurons (n = 46) showed clearly rhythmic firing during both shock-induced fast swimming and spontaneous slow swimming (Figure 7C iii). Spiking activity showed clear statistical interaction between the time of differentiation and the speed of locomotion (Figure 7D, *P* < 0.001), suggesting that groups of neurons that differed by age were recruited differently depending on the speed of locomotion. Indeed, the dorsal and middle MiV1 neurons showed significant modulation of activity based on the speed of locomotion (Figure 7D). Subthreshold activity also showed similar interaction (Figure 7E, *P* < 0.001), suggesting that this recruitment pattern can be explained at least partly by their distinct synaptic inputs. At the same time, the MiV1 group showed systematic differences in their intrinsic excitability based on their time of differentiation: the older they were, the less excitable they were (Figure 7F, *P* < 0.001). This is similar to the pattern described in the spinal cord and the caudal hindbrain V2a population (McLean et al. 2007; Kinkhabwala et al. 2011). Thus, taking advantage of whole-cell recording, we showed that these V2a reticulospinal neurons exhibited a systematic change in the spiking and subthreshold activity within each swim episode depending on the speed of locomotion. Furthermore, we revealed differential synaptic inputs and excitability linked to the birthdate-related recruitment of V2a reticulospinal neurons.

#### 4-2. Spinal innervations from individual hindbrain V2a neurons are organized based on their respective birthdates and target functionally matched spinal groups

Now that we had shown birthdate-related recruitment and its potential mechanisms in hindbrain V2a reticulospinal neurons, we examined their spinal projections in detail by labeling individual cells in each age group with dye electroporation in a series of transgenic lines labeling specific spinal neurons (Figure 8). We found distinct projection patterns across age groups at a single-cell level in a manner consisten with the overall age-related projection pattern of hindbrain V2a descending neurons, as revealed by photoconversion.

**Figure 8.**
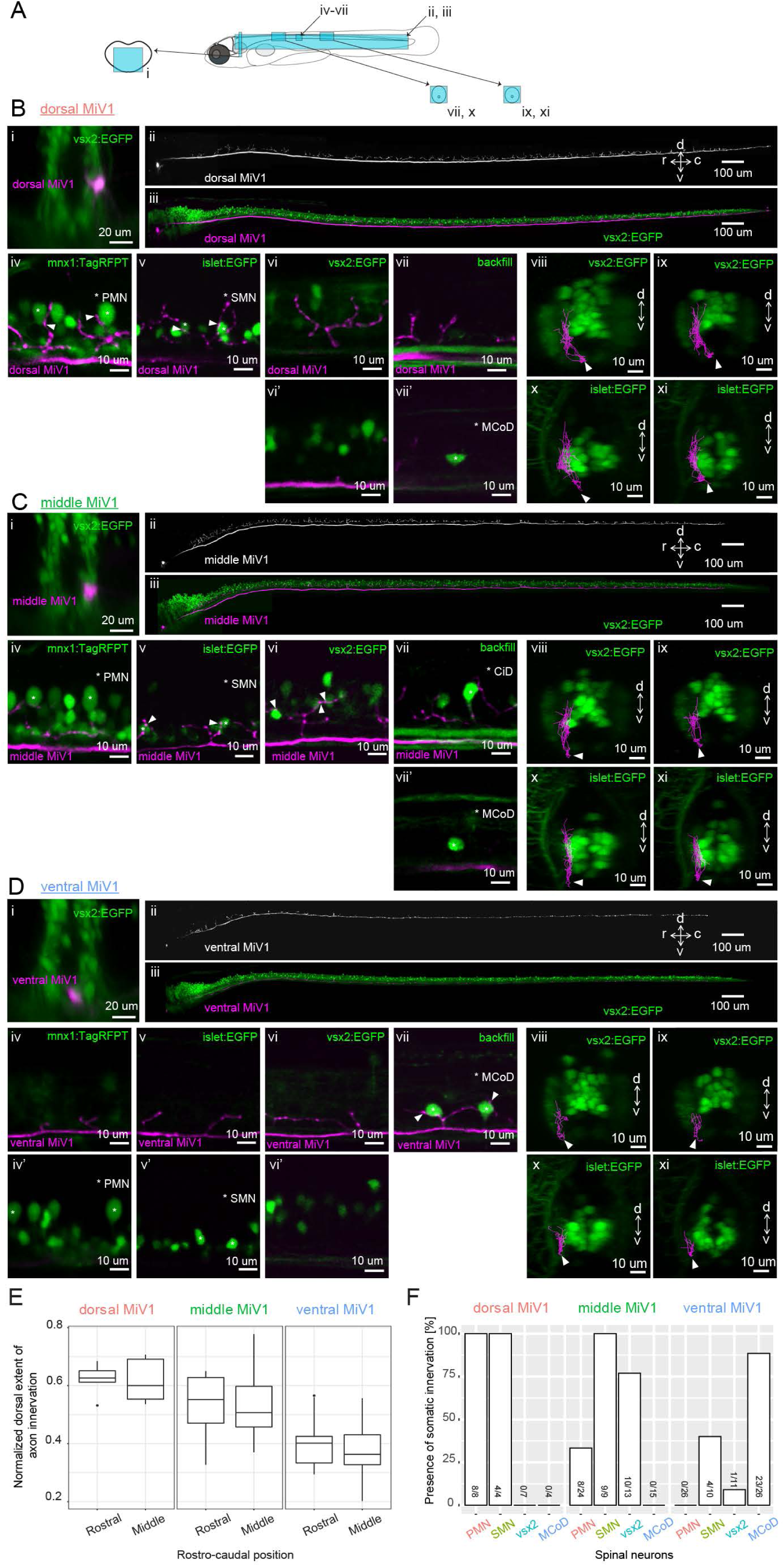
Spinal projections of V2a reticulospinal neurons in rhombomere 4. **A.** Regions displayed (cyan patches) in B, C and D. **B.** Spinal projection of an early-born V2a neuron (dorsal MiV1). (i) Coronal view of a dorsal MiV1 neuron electroporated with a dextran dye (magenta) in the background of V2a neurons (green). (ii) Side view of the spinal projection from dorsal MiV1. (iii) Same as in (ii) but overlaid on V2a neurons shown in green. (iv-vii) Sagittal optical slices showing the projection (iv) Projection relative to spinal mnx1+ neurons. Primary motoneurons (PMN) are indicated with asterisks. Processes juxtaposed to the cell body of PMN are highlighted with filled arrows. (v) Projection relative to spinal islet1+ neurons. Secondary motoneurons (SMN) are indicated with asterisks. Processes close to the cell body of SMN are highlighted with filled arrows. (vi-vi’) Projection relative to spinal V2a neurons. (vi) Optical slice showing the processes. (vi’) Optical slice medial to vi. (vii-vii’) Projection relative to a backfilled multipolar commissural descending neuron (MCoD). (vii) Optical slice showing the projection. (vii’) Optical slice lateral to vii showing the location of MCoD. (viii-xi) Coronal views of the projection of dorsal MiV1 in the rostral spinal cord (viii, x) and in the middle spinal cord (ix, xi) with V2a neurons labeled in EGFP (viii, ix) or with islet+ neurons labeled in EGFP (x, xi). **C.** Spinal projection of an intermediate V2a neuron (middle MiV1). Images are organized similarly to B. (vii) Circumferential ipsilateral descending neuron (CiD) is indicated with asterisks. **D.** Spinal projection of a late-born V2a neuron (ventral MiV1). Images are organized similarly to B. (iv-iv’) Projection relative to spinal mnx+ neurons. (iv) Optical slice showing the processes. (iv’) Optical slice medial to iv. (v-v’) Projection relative to spinal islet+ neurons. (v) Optical slice showing the processes. (v’) Optical slice medial to v. (vi-vi’) Projection relative to spinal V2a neurons, (vi) Optical slice showing the processes. (vi’) Optical slice medial to v. (vii) Projection relative to a backfilled MCoD. (vii-xi) Coronal views are organized as in B. **E.** Dorsal extent of axon innervation of V2a reticulospinal neurons in rhombomere 4. The dorsal extent of axon innervation is normalized to the thickness of spinal cord in dorsoventral axis. The main effect of hindbrain cell type (dorsal, middle, and ventral MiV1) was significant *(P* < 0.001). **F.** Somatic innervations of rhombomere 4 V2a reticulospinal neurons to spinal cell types. Percentage of cells showing somatic innervation to specific spinal cell types (PMN, SMN, vsx2, MCoD) are shown. The number of neurons showing innervation to a given class and the number of neurons examined are indicated at the bottom of each bar. The main effect of hindbrain cell type (dorsal, middle and ventral) was significant (*P* < 0.001). The interaction of hindbrain cell type and spinal cell type was also significant (*P* < 0.001).

The early-born dorsal group sent relatively thick axons through the medial longitudinal fasciculus and exhibited extensive axon collaterals throughout the spinal cord (Figure 8B ii). All the cells in this group had clear puncta-like structures - putative presynaptic terminals - in close proximity to the somata of PMNs (Figure 8B iv, 8 out of 8 cells) and SMNs (Figure 8B v, 4 out of 4 cells) but not spinal V2a neurons (Figure 8B vi, 0 out of 7 cells). Their collaterals were in the medial part of V2a neuropil (Figure 8B viii, ix) and clearly separated from the cell bodies of MCoDs located in the lateral part of the neuropil (Figure 8B vii vii’, 0 out of 4 cells).

All the cells in the intermediate age group had puncta-like structures on the ventrally located SMNs (Figure 8C v, 9 out of 9 cells) but only a third of the cells had puncta on the dorsally located PMNs (Figure 8C iv, 8 out of 24 cells). On the other hand, there was clear somatic innervation to the spinal V2a neurons in most of the cells (Fig 8C vi, 10 out of 13 cells). Again, their collaterals were in the medial part of the neuropil (Figure 8C viii, ix) and clearly separated from the laterally displaced MCoDs (Figure 8C vii, vii’, 0 out of 15 cells).

Axon collaterals from the late-born ventral group were sparse (Figure 8D ii) and located mostly in the ventral part of the neuropil (Figure 8D viii-xi). The main axon tract was in the lateral part of the neuropil where no cell bodies of motoneurons and spinal V2a neurons are present (Figure 8D iv-vi). Most of them showed clear somatic innervations to MCoDs (Figure 8D vii, 23 out of 26 cells) but far fewer or no innervations to the soma of motoneurons and spinal v2a neurons (Figure 8D iv-vi, 0 out of 26 for PMN, 4 out of 10 for SMN and 1 out of 11 for spinal V2a neuron).

At the population level, there were statistically significant differences in the dorsal extents of spinal axon arborizations across age groups (Figure 8E, *P* < 0.001). The axon arborization of each age group was also localized in a distinct medio-lateral position in the neuropil, as evident from coronal views (e.g. Figure 8B x-xi and 8D x-xi) and from their axon arborizations relative to the laterally displaced MCoDs (panels vii in Figure 8B, 8C and 8D). This is also in accordance with the chronological layering of V2a spinal projection revealed by photoconversion. These differences in axon arborization were reflected in the somatic innervation patterns to spinal neurons (Figure 8F). The interaction between MiV1 age groups and spinal cell types was statistically significant *(P* < 0.001), meaning that each age group had distinct somatic innervations based on the spinal cell types. Indeed, the dorsal early-born group was the only group that consistently innervated the cell bodies of early-born motoneurons, PMNs, whereas the ventral late-born group was the only group that had somatic innervation to late-born interneurons, MCoDs. Thus, our results provided cellular-level anatomical evidence that hindbrain V2a neurons form parallel spinal pathways arranged based on birthdate and set the stage for electrophysical analysis of these connections.

#### 4-3. Chronologically organized parallel synaptic connections to spinal neurons with distinct biophysical properties

Population- and cellular-level anatomy of hindbrain V2a neurons suggests that they exhibit distinct innervation patterns based on their time of differentiation: early-born neurons innervate early-born spinal neurons such as PMN while late-born neurons innervate late-born spinal neurons such as MCoD. To confirm these putative synaptic connectivity patterns and examine their properties in detail, we performed a series of paired whole-cell recordings from hindbrain descending neurons and spinal cord neurons (Figure 9).

**Figure 9.**
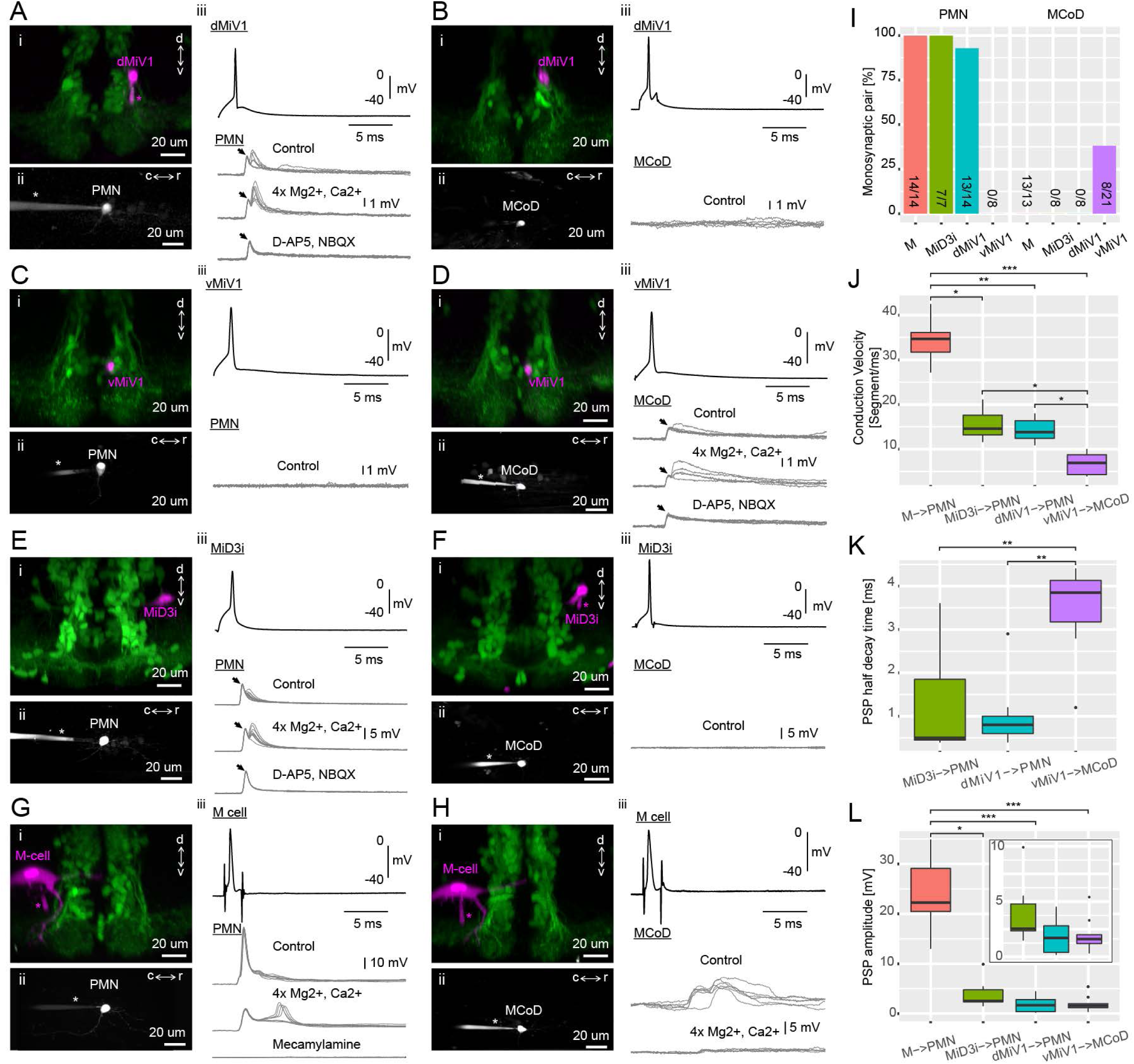
Synaptic connectivity of hindbrain V2a descending neurons. **A-H.** Paired recordings of hindbrain neurons and spinal neurons. dMiV1, dorsal MiV1; vMiV1, ventral MiV1; M, Mauthner; PMN, primary motoneuron; MCoD, multipolar commissural neuron. (i) Coronal view of a patched hindbrain neuron (magenta) overlaid on V2a neurons (green). An asterisk indicates the patch pipette in use. (ii) Side view of a patched spinal neuron (gray). (iii) Traces of postsynaptic potentials (PSPs) in a spinal neuron following spikes generated in a hindbrain neuron in control condition (top gray trace), in high divalent cation solution (middle gray trace) and following the blockade of chemical synapses (bottom gray trace). A black arrow highlights the putative electrical component of the postsynaptic potential. For simplicity, only one action potential trace is shown (top black trace). **I.** Monosynaptic connections of hindbrain descending neurons to spinal neurons. Percentage of pairs showing monosynaptic connections to spinal neurons are shown. The number of connected pairs and the number of pairs examined are indicated at the bottom of each bar. The main effect of hindbrain cell types on the percentage of monosynaptic connection was significant (*P* < 0.001). The main effect of spinal cell types was also significant (*P* < 0.001). **J.** Conduction velocity of monosynaptically connected pairs. Conduction velocity was defined as the number of muscle segments an action potential propagated in a millisecond. Box and whisker plots represent median as well as first and third quartiles. Asterisks mark significant differences between given connections (Dunn’s test, \**P* < 0.05, \*\**P* < 0.01, \*\*\**P* < 0.001). **K.** Half decay time of monosynaptic PSPs. Data are presented as in J. **L.** Amplitude of monosynaptic PSPs. Data are presented as J. (inset) Zoomed-in box and whisker plots for the pairs with smaller PSP.

First, we examined the synaptic connectivity of dorsal MiV1, the early-born V2a descending neurons in rhombomere 4, to PMN (Figure 9A; n = 14) and MCoD (Figure 9B; n = 8). The firing of an action potential in dorsal MiV1 led to a postsynaptic potential in PMN in most pairs (Figure 10A; 13 out of 14). In 8 cases, the EPSPs had two clear components: a short latency component and a relatively longer latency component (Figure 9A iii). We tested the monosynaptic nature of the connections by raising magnesium and calcium concentrations to minimize the contribution of polysynaptic pathways (Figure 9A iii). In these cases (n = 7), the responses persisted, suggesting that they were mediated by monosynaptic connections. The longer latency component was blocked by a mixture of glutamatergic blockers, leaving only the shorter latency component (Figure 9A iii; n = 7). The longer latency component was also variable across trials while the short latency component was stable across trials. Therefore, the initial short latency response was probably due to an electrical synapse, and the later response that followed this was due to a glutamatergic synapse. On the other hand, none of the MCoDs showed a postsynaptic response to an action potential elicited in the dorsal MiV1 (Figure 9B; 8 out of 8). Taken together, our results indicate that in a manner consistent with their spinal projection patterns, dorsal MiV1s, the early-born V2a neurons in rhombomere 4, show selective synaptic connectivity to PMNs, the early-born spinal neurons.

**Figure 10.**
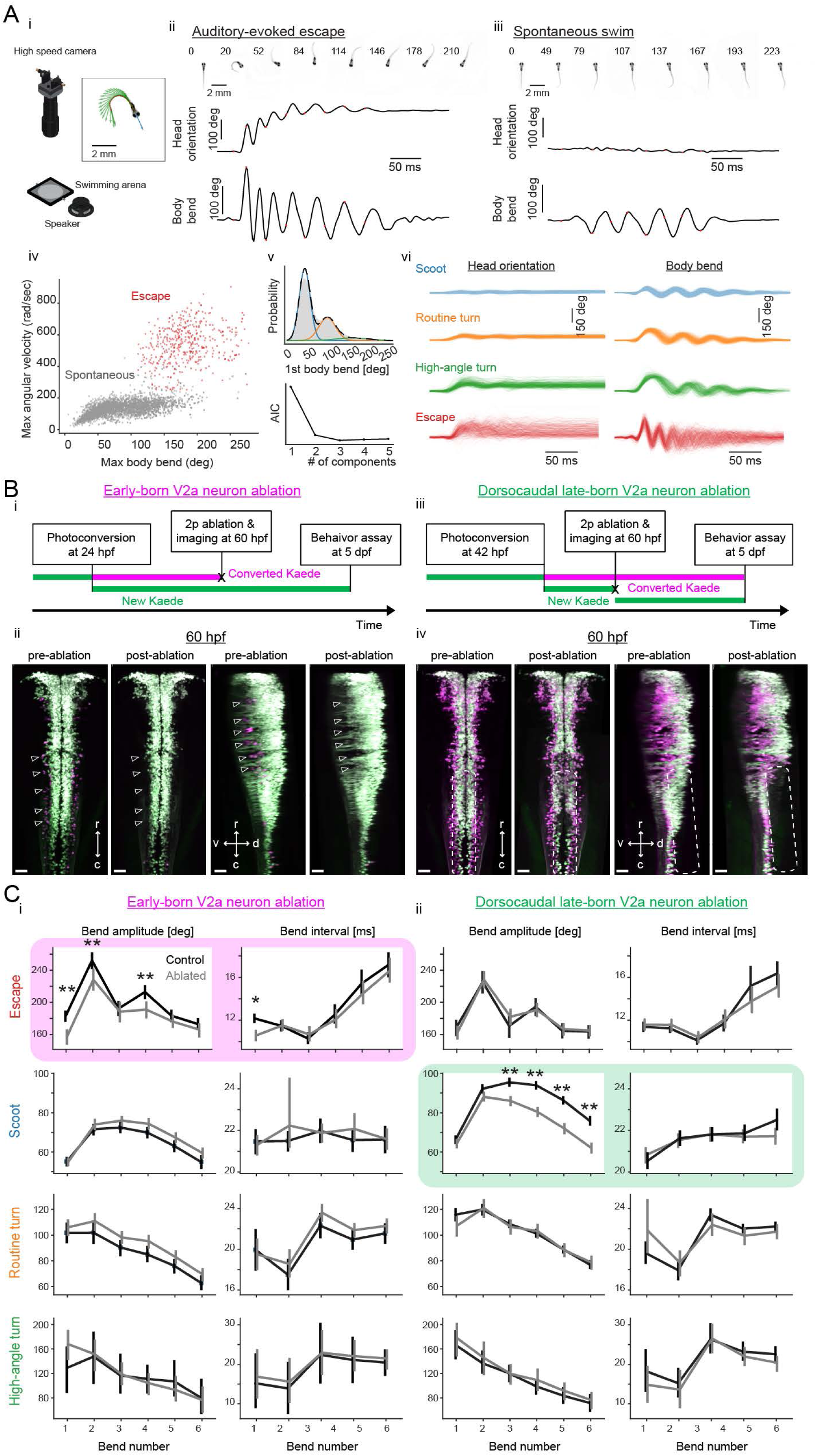
Contributions of hindbrain V2a neurons to evoked and spontaneous locomotion. **A.** Examined locomotion patterns. (i) Experimental setup. (inset) Example fish image showing tracking of head orientation and tail curvature (see Methods). Blue arrow, head orientation; Red dots, tracked midline points; Green arrows, tangent vectors along the midline points. (ii) Auditory-evoked escape response. (Top row) Images of fish during escape response. The number on each the images is time from the stimulus onset in milliseconds. (Middle row) Time course of head orientation. (Bottom row) Time course of total body bend (see Methods). (iii) Spontaneous swimming. Panels are organized similarly to ii but the number on each image is time from the movement onset in milliseconds. (iv) Scatter plot of maximum bend amplitude (during a swim episode) and maximum angular velocity for auditory-evoked escapes (n = 426 episodes, N = 10 fish, red points) and spontaneous swims (n = 6088 episodes, N= 15 fish, gray points). (v) (Top) Probability of initial bend amplitude for 6056 spontaneous swim events. A fit of gaussian mixture model identified three components. (Bottom) The Akaike information criterion (AIC) as a function of number of components. Three components led to the lowest AIC value. (vi) Overlaid traces of head orientation and total body bend for three identified spontaneous swim categories (Scoot, 3969 episodes, blue; Routine turn, 1565 episodes, orange; High-angle turn, 376 episodes, green) and auditory-evoked escape (Escapes, 381 episodes, red). **B.** Femtosecond laser ablation of hindbrain V2a neurons. (i) Experimental procedure for the ablation of early-born V2a neurons. (ii) Maximum intensity projections before and after ablation at 60 hpf (left, dorsal view; right, side view). White open arrowheads indicate the locations of the early-born V2a neurons.(iii) Experimental procedure for the ablation of dorsocaudal late-born V2a neurons. (iv) Maximum intensity projections before and after ablation at 60 hpf (left, dorsal view; right, side view). White rounded rectangles indicate the location of the late-born dorsocaudal V2a neurons, r, rostral; c, caudal; d, dorsal; v ventral; scale bars, 30 um. **C.** Effects of hindbrain V2a neuron ablations on four distinct locomotion patterns. (i) Effects of the early-born V2a neuron ablation. Bend amplitude and interval are quantified bend-by-bend from 1^st^ bend to 6^th^ bend (see Methods). Asterisks mark significant differences between ablated and control fish (** p< 0.01, corrected for multiple comparisons). Error bars, 99 percent confidence interval. The swim category that showed significant effects are highlighted in magenta. (ii) Effects of the late-born dorsocaudal V2a neuron ablation. Panels are organized as in i. The swim category that showed significant effects are highlighted in green.

**Figure 11.**
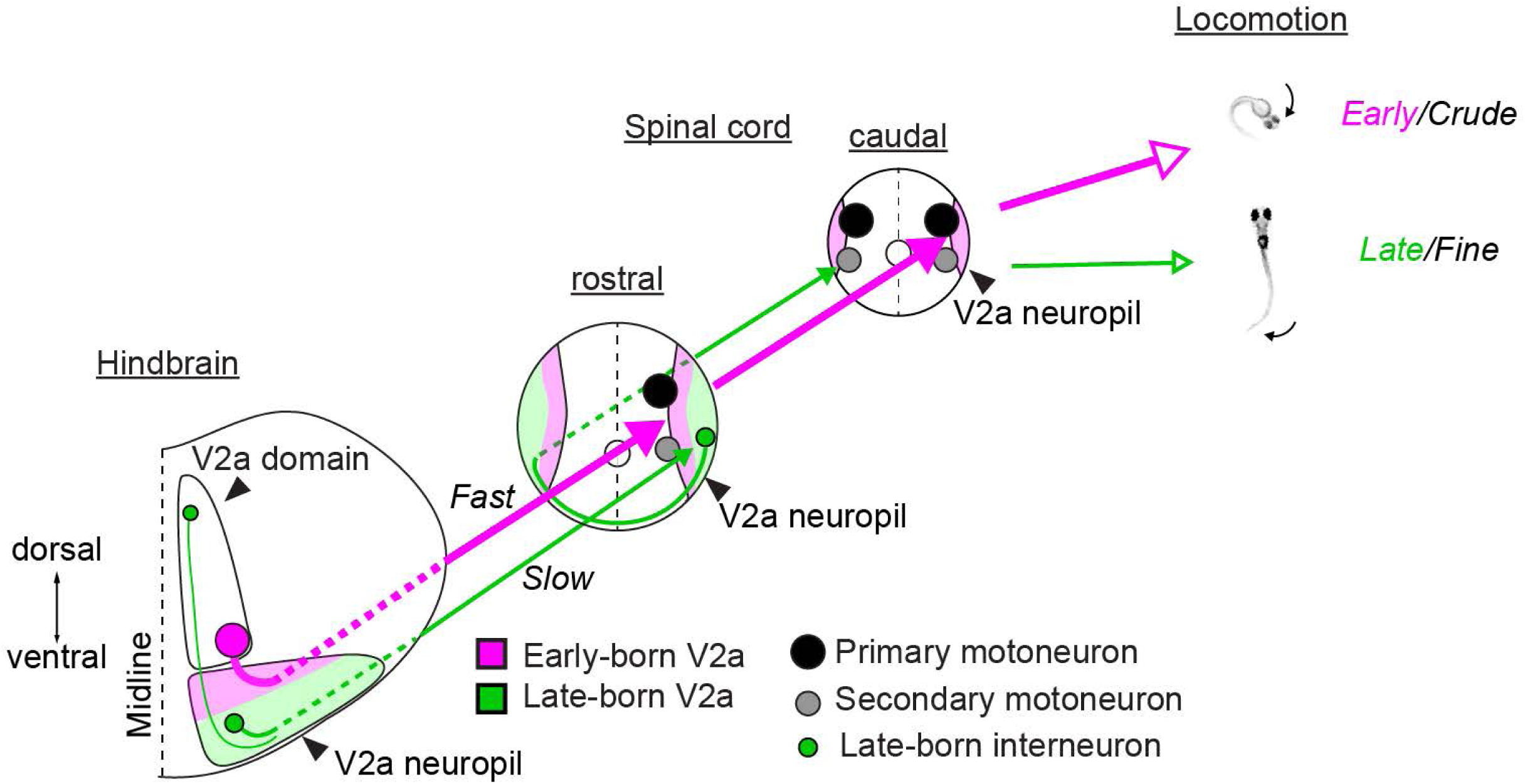
Chronological architecture of hindbrain V2a descending pathways that underlies diversification and increase in sophistication of zebrafish locomotor behaviors. A schema illustrating the relationship between the birthdate-related organization of hindbrain V2a descending pathways and the development of locomotor behaviors. Cell bodies are represented in circles. A larger circle means lower excitability. Arrows indicate descending axons. A larger arrow indicates faster and stronger connection. Neuropil regions are color-coded to indicate the separation based on the time of differentiation. Magenta area, early-born neuropil; Green area, late-born neuropil. Open arrows indicate the correspondence between pathways and axial movements during locomotion. The size of open arrow indicates the strength of the movement.

In contrast, ventral MiV1s, the late-born V2a descending neurons in rhombomere 4, showed the opposite pattern of synaptic connectivity. None of the PMNs showed a postsynaptic response to an action potential elicited in ventral MiV1 (Figure 9C; 0 out of 8) while one third of MCoDs examined showed a clear postsynaptic potential (Figure 9D; 8 out of 21). The responses in these MCoDs persisted in high magnesium and calcium solution, suggesting they were generated monosynaptically. In four out of eight cases, we observed clear biphasic responses (Figure 9D iii), the latter component of which was blocked by glutamatergic blockers (Figure 9D iii; n = 4). The amplitude of the short latency component was stable across trials in contrast to the later component mediated by glutamatergic synapses, suggesting that the early component was probably due to electrical synapses. Thus, in a manner consistent with their spinal projection patterns, ventral MiV1s showed selective monosynaptic connectivity to MCoDs, the late-born spinal interneurons that are only active during slow swimming (McLean et al. 2008).

To see if other V2a descending neurons and non-V2a descending neurons show similar connectivity patterns based on time of differentiation, we also looked at the connectivity of early-born V2a neurons in rhombomere 6, MiD3i, (Figure 9E, 10F) and the earliest born descending neurons in rhombomere 4, the Mauthner cell (M-cell), (Figure 9G, H) to PMNs and MCoDs. In 7 out of 7 cases, a clear biphasic postsynaptic potential was observed in PMNs in response to an action potential in MiD3i (Figure 9E). This response persisted in high magnesium and calcium solution, suggesting that they were monosynaptic. Once again, the early component was stable across trials while the late component was variable and blocked by glutamatergic blockers (Figure 9E iii; n = 7), suggesting that they were electrical and glutamatergic respectively. The M-cell also showed similar, but much stronger connections. In 14 out 14 cases, PMNs showed very strong postsynaptic potentials leading to spiking activity in response to a single action potential in the M-cell (Figure 9G iii). These responses also persisted in high magnesium and calcium solution, suggesting that the connections were monosynaptic. These responses were completely blocked by a cholinergic blocker (Figure 9G iii; n = 10), indicating that they were mediated by cholinergic synapses, in accordance with existing literature (Koyama et al. 2011; Clemente et al. 2004). MCoDs on the other hand showed a long latency complex postsynaptic potential in response to one M-cell action potential, suggesting that the connection is probably polysynaptic (Figure 9H iii; 13 out of 13 cases). Most, if not all, of the responses disappeared in high magnesium and calcium solution, further supporting that they are mediated polysynaptically (Figure 9H iii; n = 10). In sum, our findings indicate that both types of early-born descending neurons make synaptic connections to the early-born PMNs but not to the late-born MCoDs.

Overall, the connection probability was statistically different depending on the hindbrain cell type and the spinal cord cell type (Figure 9I; main effect of hindbrain cell type, *P* < 0.001; main effect of spinal cell type, *P* < 0.0001). In fact, PMNs and MCoDs received synaptic inputs from completely different sets of reticulospinal neurons, with each set corresponding to distinct age group (Figure 9I). Thus, these results showed electrophysiologically that early-born and late-born hindbrain neurons form parallel pathways to the spinal cord and these pathways are organized based on the time of differentiation, supporting the idea that parallel hindbrain-spinal cord circuits emerge in sequence during development.

We then examined biophysical properties of these synaptic connections. We first examined differences in conduction velocity among connected pairs (Figure 9J): the youngest ventral MiV1/MCoD pairs were the slowest among the connected pairs. The middle-aged pairs (dorsal MiV1/PMN and Mid3i/PMN) were slower than the oldest M-cell/PMN pairs. We also examined differences in the time constants of synaptic potentials, excluding the M-cell/PMN pair that showed spiking activity (Figure 9K). The youngest ventral MiV1/MCoD pairs showed significantly longer time constants than the older V2a neurons (Figure 9K). With respect to the amplitudes of the synaptic potentials, the M-cell/PMN pair was the only pair that was statistically different from other pairs (Figure 9L). However, consistent with previous studies (McLean et al. 2007; Menelaou and McLean 2012), the early-born PMNs and the late-born MCoDs examined here differed significantly in input resistance (92.3±18.2 M Ohm and 514±172 M Ohm, respectively; Wilcoxon test, *P* < 0.001), suggesting that the underlying synaptic currents were systematically stronger for the early-born hindbrain neurons than the late-born hindbrain neurons. Altogether, we found systematic differences in conduction velocities, synaptic decay times and synaptic currents between the two parallel pathways: the early-born pathway is equipped with fast conduction velocities, fast time constants of synaptic potentials and stronger synaptic currents while the late-born pathway is equipped with slow conduction velocities, slow time constants of synaptic potentials and weaker synaptic currents. The differences in the biophysical properties as well as connectivity patterns are consistent with the distinct types of axial motor activity observed during the locomotor patterns in which these pathways participate. The net effects of the distinct biophysical properties and connectivity patterns associated with the early- and late-born pathways is that the former could more quickly bring the PMNs throughout the spinal cord to their firing thresholds, whereas the latter could more slowly bring the SMNs in the caudal spinal cord to their firing thresholds. As a consequence, activation of early-born pathways could produce more powerful whole body bending observed during different crude forms of locomotion (McLean et al. 2008; Liao and Fetcho 2008), whereas activation of late-born pathways could produce weaker bending of the caudal tail region with a long rostrocaudal delay, as observed during the refined locomotion (McLean et al. 2008). In sum, these results suggest that the parallel hindbrain-spinal cord circuits that develop in sequence and have distinct connectivity patterns and biophysical properties contribute to the distinct locomotor patterns that zebrafish display in sequence.

### 5. Behavioral contributions of distinct age groups of hindbrain V2a descending neurons

Systematic characterization of the development, recruitment, and connectivity of V2a descending neurons led us to hypothesize that early-born hindbrain V2a descending neurons contribute to the stimulus-evoked crude locomotion that develops early in development while their late-born counterparts contribute to the more refined spontaneous locomotion that develops later. To test this hypothesis, we separately ablated either the early-born or the late-born hindbrain V2a descending neurons and at 5 days post-fertilization (dpf) examined the effects of each of these ablations on a few distinct patterns of locomotion (Figure 10).

To examine a wide range of swimming patterns in the freely swimming condition, we employed two experimental conditions: auditory-evoked escape responses (Figure 10A ii) and spontaneous swimming (Figure 10A iii). In each condition we monitored movements of the fish using a high-speed camera (Figure 10A i). Auditory stimulus-elicited escape episodes consisted of large and fast changes in head orientation as well as body bends (Figure 10A ii), these being characteristic of the crude locomotion that develops early. On the other hand, spontaneous swim episodes mostly consisted of small changes in head orientation, and slow and small bends restricted to the caudal portion of the tail (Figure 10A iii), and these are typical of the more refined locomotion that develops later. The distinctiveness of these two swim types became quite apparent when for each swim episode the maximum angular velocity (see Methods) was plotted against the maximum body bend amplitude (Figure 10A iv). In this plot, the points corresponding to escapes and slow swims clearly segregated into distinct clusters. Consistent with the findings of a previous study (Burgess and Granato, 2007), when we looked at the distribution of the amplitude of the first total body bend in each spontaneous swim episode we found it to be multimodal, suggesting that spontaneous swim episodes consisted of distinct subtypes (Figure 10A v). A Gaussian mixture model fit to the distribution suggested that there were potentially three subcategories of spontaneous swim episodes (see Methods). The timeseries of body bends and head orientation in each category showed consistent motor patterns (Fig 10A vi), further supporting the idea that fish at 5 dpf exhibit at least three distinct motor patterns spontaneously. We referred to these patterns as “scoot”, “regular turn” and “high-angle turn” based on the nomenclature adopted in previous studies (Burgess and Granato 2007; Marques et al. 2018). Combined with auditory-evoked escape responses, we examined four distinct motor patterns in total (Figure 10A vi).

We focused on two age groups of hindbrain V2a descending neurons based on their times of differentiation and their recruitment patterns. The first group consisted of neurons born before 24 hpf (10 fish for the ablated group; 10 fish for the control group). We targeted all of these neurons in the hindbrain as the optical backfill and Ca^2+^ imaging indicated that all of them were descending neurons involved in strong locomotion. We visualized these neurons by photoconverting Tg(vsx2:Kaede) at 24 hpf, and at 60 hpf we ablated these neurons by femtosecond laser pulses (Figure 10B i; 131 ± 17.3 cells/fish). We delivered multiple pulses to each neuron (typically 5-10 pulses/neuron) until the fluorescence signal suddenly decreased. Neurons labeled with photoconverted Kaede disappeared with no clear unintended damage to the nearby cells and processes (Figure 10B ii). To confirm that these neurons did not recover before the behavior experiments, we imaged them again at 4 dpf (Figure 10 supplement 1A). Comparison with the control group at 4 dpf clearly showed that these early-born neurons did not recover from ablation. The second group of neurons targeted for ablation consisted of neurons born from 42 hpf to 60 hpf (Figure 10B iii; 10 fish for the ablated group; 10 fish for the control group). We focused on the neurons in the dorsocaudal hindbrain because the optical backfill and Ca^2+^ imaging indicated these neurons are descending neurons involved in spontaneous slow locomotion. We identified these V2a neurons based on the lack of photoconverted Kaede at 60 hpf after the photoconversion at 42 hpf (Figure 10B iii) and their relative position to the nearby early-born neurons (508 ± 15 cells/fish; see Methods). Femtosecond laser pulses ablated these neurons with no clear unintended damage to the nearby early-born neurons (Figure 10B iv). Comparison with the control group at 4 dpf clearly showed the reduction of the late-born neurons in the dorsocaudal hindbrain (Figure 10 supplement 1B).

We examined the kinematic differences in axial movements between the ablated and control group for each age group. We did this separately for each of the four distinct motor patterns. Specifically, we examined the amplitude and the duration of each body bend (Figure 10C), but only in the part of the episode (initial six bends) that is reliably observed across episodes (Figure 10A vi). Relative to the control group, the early-born V2a ablation group exhibited significant decreases in the amplitudes of the first, second and fourth bend, as well as a decrease in the duration of the first bend of escape (Figure 10C i, Escape; *P* < 0.01, corrected for multiple comparisons). However, we observed no statistically significant changes in the three other slower locomotor patterns (Figure 10C i, Scoot, Regular turn and High-angle turn). On the other hand, in the late-born V2a ablation group we saw no clear changes in the escape response (Figure 10C ii, Escape). Instead, this group showed a decrease in the amplitudes of bends in the later phase of scoot (Figure 10C ii, Scoot; *P* < 0.01, corrected for multiple comparisons) but not the other two spontaneous swim patterns involving turning (Figure 10C ii, Regular turn and High-angle turn). These observations comport well with the recruitment pattern and the spinal projections we observed earlier of the neurons we ablated here. Furthermore, the fact that we observed significant changes only in the late phase of scoots suggests that late-born V2a neurons may play a role in maintaining the excitatory drive in the spinal cord during slow forward locomotion.

To gain assurance that we compared the same set of spontaneous locomotor behaviors between the control groups and the ablated groups, we compared the distributions of the first body bend we used to subcategorize spontaneous locomotor behavior (Figure 10 supplement 2). Although there were slight differences in the peak amplitudes of the probability distributions across the groups, the overall shapes of the distributions were consistent, indicating that we compared the same sets of locomotor behaviors across the control and ablated groups. To examine this more closely for scoot behavior, which showed significant deficits upon ablation of late-born caudal hindbrain neurons, we examined the time course of the amplitude of body bending during this behavior (Figure 10 supplement 3). Traces from all the episodes of scoot from the ablated fish (Figure 10 supplement 3, orange) and the control fish (Figure 10 supplement 3, blue) are overlaid for each ablation group. In both ablation groups, the ablated and control groups both exhibited the small amplitude body bends with slow left-right alternation (Figure 10 supplement 3) that is characteristic of scoot behavior (Figure 10 A vi). Furthermore, the overlaid traces of the late-born V2a ablation and control groups clearly revealed that the bend amplitudes of scoots decreased in the later phase of scoot in the ablated group (Figure 10 supplement 3A). This further strengthened the idea that late-born V2a neurons of the caudal hindbrain play a role in the maintenance of excitatory drive during slow forward locomotion. To summarize, our ablations did not change the set of distinct locomotor behaviors fish exhibit spontaneously.

While examining the results of ablations we noticed that the early-born and the late-born control groups exhibited significantly different bend amplitudes during scoot (Figure 10C i, Scoot; Figure 10C ii, Scoot, *P* < 0.01). Comparison to an independent control group that was not subjected to photoconversion suggested that photoconversion at 42 hpf impacted the kinematics of scoot at 5 dpf (data not shown). Although this does not undermine the observed significant decrease in bend amplitudes of scoot in the late-born ablation group, it does raise some concerns about the lack of significant effects in the early-born ablation group. For instance, it is conceivable that ablation of early-born neurons does not reveal a decrease in scoot bend amplitudes because of a floor effect, wherein the observed bend amplitudes somehow represent the lowest values that the fish can inherently display in order to produce a scoot (Fig. 10C i, Scoot). However, we think this is unlikely because we noticed a slight increase rather than decrease in bend amplitudes after ablation of the early-born neurons (Figure 10C i, Scoot). In any case, the effects of photoconversion itself further justifies our use of matched control for each ablation group.

In conclusion, with ablation experiments we confirmed our hypothesis about the functional roles of early-born and late-born V2a neurons in freely swimming fish. Moreover, the double dissociation of the two locomotor patterns indicates that the underlying V2a descending pathways are largely independent, in accordance with their parallel circuit arrangement resulting from the layered growth of their spinal projections. This suggests that the layered development of new connections in the nervous system underlies its ability to integrate new circuits into existing circuitry in such a way as to allow the different circuits to operate autonomously. Interestingly, previous studies showed that the ablation of ventral reticulospinal neurons, the hindbrain V2a neurons we classified as the intermediate-aged group, affects slow but large turns with no clear kinematic effects on forward locomotion and suggested that the descending pathway for turning behaviors is independent of the one for forward locomotion (Huang et al. 2013; Orger et al. 2008). Collectively, this suggests that the array of locomotor behaviors fish display after birth are supported by sequentially developing parallel hindbrain-spinal cord circuits, with each circuit accounting for a specific range of kinematics and dynamics through its unique connectivity pattern and biophysical properties.

## Discussion

We examined hindbrain V2a neurons and their descending pathways and uncovered a chronology-based architecture that explains the postnatal development of locomotor behaviors (Figure 12). Hindbrain V2a descending neurons born early are also the first to project to the spinal cord where they innervate similarly early-born motoneurons throughout the spinal cord with fast conducting axons and synapses that exhibit fast dynamics. Consistent with their connectivity pattern and biophysical properties, they contribute to the crude and fast locomotion that consists of fast and powerful bends of the whole-body. Hindbrain V2a descending neurons born later give rise to a more laterally located layer of descending pathways that function in parallel to existing pathways: through similarly late-born premotor neurons in the rostral spinal cord, these V2a neurons polysynaptically connect to caudal motoneurons via slow conducting axons and synapses that exhibit slow dynamics. Consistent with the connectivity pattern and biophysical properties, they contribute to the more refined and slower locomotion that appears later and that consists of slower and weaker bends of the caudal tail. In sum, our results reveal that the chronological layering of parallel circuits and the systematic variation in the connectivity patterns and biophysical properties of these circuits underlie the diversification and increase in sophistication of the locomotor repertoire.

Such chronology-based incorporation of new motor circuits in parallel to existing circuits may provide distinct advantages over non-parallel organization such as exists within the midbrain descending pathways (Wang and McLean 2014). First, it allows the motor patterns produced by these circuits to be controlled independently of one another. Such independent control may be critical for some behaviors such as prey capture, for instance, that require flexible coordination of multiple motor patterns. Indeed, during the final approaching phase of prey capture, a zebrafish can increase the speed of forward swimming to values approaching those seen during fast behaviors established earlier in development, but without displaying the large lateral head displacements seen in these earlier-established behaviors (Patterson et al. 2013). This suggest that a fish can generate fast forward swimming by strongly driving only the late-born pathway that predominantly drives tail movements. Second, this parallel organization allows incorporation of new circuits controlling increasingly sophisticated motor patterns in an open-ended way. Interestingly, the development of the corticospinal pathway in mammals appears to fit into this view; in this pathway spinal tracts are established postnatally in the lateral part of the spinal cord after the prenatal development of other descending pathways (Lakke 1997; Kuypers 1962), and these tracts contribute to skilled forelimb movements that animals gradually develop postnatally (Martin 2005; Alstermark and Isa 2012). Further analysis will be necessary, however, to assess the degree to which the corticospinal pathway can function without contributions from other descending pathways (see Esposito, Capelli, and Arber 2014). In sum, chronology-based parallel incorporation of new motor circuits provides certain advantages over non-parallel organization.

There is some evidence that suggests that such chronology-based parallel incorporation of new circuits extends beyond motor systems. In the lateral line system of zebrafish, sensory afferents in the hindbrain are topologically organized based on their birthdate (Pujol-Martí et al. 2012). In the cerebellar molecular layer in mice, parallel fiber axons from granule cells line up in order of age (Espinosa and Luo 2008). In the fly visual system, the growth cones of photoreceptor afferents are segregated based on their birth order (Kulkarni et al. 2016). Also, in the mouse hippocampus, age-matched subpopulations of principal neurons are interconnected (Deguchi et al. 2011). Although the link between age-dependent layered circuit organization and the development of behaviors remains to be demonstrated in these systems, given how widely this pattern is observed, we suspect that it is a fundamental organizing principle in the brain that allows for implementation of new circuits in parallel to existing ones to meet the demands of an increasingly diverse behavioral repertoire following an animal’s birth.

The systematic changes in connectivity patterns and biophysical properties we observed in the chronologically layered parallel circuits fit well with the general development pattern of movements observed in many vertebrates (Kagan and Herschkowitz 2006; Harlow and Harlow 1965; Fox 1965; Drapeau et al. 2002). For example, in humans, global and fast body movements such as startle develop first followed by isolated limb movements and fine motor skills (de Vries, Visser, and Prechtl 1982; Gallahue, Ozmun, and Goodway 2012). Broad and direct connectivity patterns equipped with fast biophysical properties would be suitable for the global and fast body movements that develop early. Localized and indirect connectivity patterns equipped with slow biophysical properties would be suitable for the finer movements that develop later. It would be interesting to see if the neural pathways underlying such behaviors indeed exhibit systematic changes in connectivity patterns and biophysical properties.

The aforementioned systematic changes in biophysical properties and connectivity patterns may not be restricted to motor systems. Indeed, correlations between morphological and physiological properties and the time of differentiation have also been reported in the mammalian cortex (Butt et al. 2005; Miyoshi et al. 2007) and hippocampus (Picardo et al. 2011; Marissal et al. 2012). Although the computational and behavioral roles of these systematic changes in morphological and physiological properties remains to be answered, it seems likely that they represent a general strategy utilized by nervous systems to support the wide range of neural processes required for the complete behavioral repertoire of animals: from the fast and global sensorimotor processing observed in innate reflexive behaviors to the more sustained and finer processing presumed to happen in deliberate and sophisticated cognitive behaviors that animals acquire later.

The hindbrain contains a series of cell types that are arranged in columns that run rostrocaudally throughout the hindbrain (Kinkhabwala et al. 2011; Koyama et al. 2011; Gray 2013). Each cell type exhibits a specific combination of transcription factor expression, neurotransmitter identity and axonal projection pattern. Hindbrain V2a neurons comprise one such cell type. As these cell types span across various sensorimotor circuits in the hindbrain, it has been proposed that each cell type may provide a fundamental neural operation that is broadly useful to many of these circuits (Kinkhabwala et al. 2011; Koyama and Pujala 2018). Indeed, there is some evidence suggestive of the involvement of hindbrain V2a neurons in other hindbrain motor circuits besides locomotor circuits. In mice, genetic ablation of all V2a neurons in the nervous system produces deficits in respiration (Crone et al. 2012). A class of premotor neurons in the horizontal eye movement circuit partly corresponds to putative V2a neurons in larval zebrafish (Lee, Arrenberg, and Aksay 2015). These neurons show persistent activity correlated to eye position but the time constant of persistent activity varies across cells (Miri, Daie, Arrenberg, et al. 2011). This raises the possibility that, as in locomotor movements, eye movements are controlled by a series of V2a neurons that show systematic changes in biophysical properties based on the time of differentiation. Perhaps one fundamental role of hindbrain V2a neurons is to provide to hindbrain motor circuits a series of excitatory drives that vary in duration so that these circuits can result in a behavioral repertoire containing movements with a wide range of temporal dynamics. Precise functional mappings and detailed circuit analyses will be necessary to examine if this is indeed the case.

Although the parallel arrangement of differentially-aged circuits that control distinct behaviors can explain how these behaviors can be controlled independent of one another, it does not explain how animals can coherently string together these behaviors. The coordination between early-born and late-born neurons has been described in a few neural systems. For example, through their widespread axonal arborizations, early-born “pioneer” GABAergic neurons in the hippocampus exert a huge impact on the activity of neurons born afterwards, and in this way these neurons play a crucial role in the synchronization of activity observed in the developing hippocampus (Picardo et al. 2011). Although these early-born neurons have been shown to exist in the adult hippocampus and to project heavily to the septum (Villette et al. 2016), it has yet to be revealed how their control over late-born neurons influences behavior in adulthood. The reticulospinal system in the larval zebrafish could serve as a good avenue for understanding the interactions that occur among age groups during behavior because it is composed of a relatively small number of identifiable neurons, many of which are known to be recruited during the same behavior (Kimmel, Powell, and Metcalfe 1982; Gahtan et al. 2002). This may permit detailed analyses of the interactions among age groups and their behavioral roles. In any case, detailed circuit analyses of early-born and late-born neurons in the context of behavioral development will be essential to understand how nervous systems organize the interactions between early-born and late-born circuits, and thus allow animals to produce coherent behaviors.

## Materials and methods

### Fish care

Zebrafish larvae were obtained from an in-house breeding colony of wild-type adults maintained at 28.5 °C on a 14-10 hour light-dark cycle. Embryos were raised in a separate incubator but at the same temperature and on the same light-dark cycle. Embryos were staged (Kimmel et al. 1995) and only the ones with normal development were used in the study. All experiments presented in this study were conducted in accordance with the animal research guidelines from the National Institutes of Health and were approved by the Institutional Animal Care and Use Committee and Institutional Biosafety Committee of Janelia Research Campus.

### Transgenic fish

The following previously published transgenic lines were propagated to Casper background and used in this study: TgBAC(vsx2:GFP)(Kimura, Okamura, and Higashijima 2006); TgBAC(vsx2:Kaede)(Kimura, Okamura, and Higashijima 2006); TgBAC(vsx2:Gal4)(Kimura et al. 2013); Tg(UAS:Kaede)(Davison et al. 2007); Tg(UAS:synaptophysin-EGFP)(Heap et al. 2013); Tg(UAS:GCaMP6s) (Muto et al. 2017); Tg(mnx1:TagRFPT)(Jao, Appel, and Wente 2012); TgBAC(islet1:GFP)(Higashijima, Hotta, and Okamoto 2000); Tg(Dbx1b:Cre, vglut2a:lRl-GFP)(Koyama et al. 2011). For examining cholinergic transmission in the Mauthner cell, we used the paralytic mutant, *relaxed*, to avoid the use of the cholinergic blocker α-bungarotoxin for the induction of paralysis (Koyama et al. 2011).

### Developmental imaging of hindbrain V2a neurons

TgBAC(vsx2:EGFP) was anesthetized in tricaine methanesulfonate (MilliporeSigma, E10521, St Louis, MO) dissolved in system water at 160 mg/L (hereafter referred as MS-222) and then embedded in 1.6% low melting point agar (MilliporeSigma, 2070-OP, MO). Volumetric images of hindbrain V2a neurons were acquired with a custom two-photon microscope. 25x 1.1 NA objective lens (Nikon Instruments, CFI Apo LWD 25XW 1300nm, NY) was used and EGFP was excited at 940 nm (MKS Instruments, Mai Tai HP, Deep See, MA). For each developmental time point, a separate group of fish was used to avoid the potential developmental effects of the imaging procedure ( > 6 fish per time point). For time lapse imaging, fish were kept in 0.5 % agar and temperature was maintained at 28.5 °C with a temperature controller (Luigs & Neumann, TC07, Germany).

### Birthdating of hindbrain V2a neurons using a photoconvertible fluorescent protein

Fish expressing the photoconvertible fluorescent protein, Kaede, in V2a neurons (TgBAC(vsx2:Kaede)) were photoconverted at 24, 36, 48, 60, 72 and 96 hours post fertilization (hpf) to identify V2a neurons that existed at a given developmental time point using a procedure similar to the one described previously (Kimura, Okamura, and Higashijima 2006; Caron et al. 2008). Briefly, UV light was shone on around 10 fish within a drop of system water for 30-60 sec using a stereo dissection microscope (Olympus, MVX10, PA) equipped with a DAPI filter (Chroma Technology, 49901, VT) and a halogen bulb (Excelitas Technologies, X-Cite 120PC Q, MA). Then, the fish were transferred back into the incubator and housed in a box that filtered out UV wavelengths from the light within the incubator. This prevented photoconversion of additional developing neurons by the ambient light in the incubator. The fish remained in the incubator until 120 hpf, at which point they were imaged with a confocal microscope (Zeiss, LSM710, Germany) with 20x 1.0 NA objective lens (Zeiss, W Plan-Apochromat 20x/1.0, Germany) (> 8 fish per time point). A group of fish converted at 24 and 36 hpf (6 fish per time point) were injected with a dye in the spinal cord (Liu and Fetcho 1999) at 4 days post fertilization (dpf) to label reticulospinal neurons. A far red dye was used to avoid contaminating the red channel used for imaging photoconverted Kaede (Thermo Fisher Scientific, Alexa Fluor 680 dextran, MW: 10,000, MA). These neurons are identifiable across animals and are named based on their anatomical features (Kimmel, Powell, and Metcalfe 1982; Mendelson 1986). We used the same naming convention used previously except for a V2a reticulospinal neuron that was born by 24 hpf in rhombomere 4. This cell was the most dorsal cell within V2a reticulospinal neurons in rhombomere 4 but was slightly ventral to MiM1. Just as MiM1 refers to a single identifiable cell that is just ventromedial to the Mauthner cell (Mendelson 1986), we decided to refer to this neuron as dorsal MiV1.

### Optical backfill of hindbrain V2a neurons

Fish expressing Kaede in V2a neurons through the Gal4-UAS system were photoconverted in the rostral spinal cord at 36, 60 and 84 hpf to identify hindbrain V2a neurons that had spinal projections at a given developmental time point as described previously (Kimura et al. 2013). Briefly, fish were anesthetized in MS-222 and embedded ventral-side up in 0.5% low-melting point agar and then the rostral spinal cord (muscle segment 5 to 7) was illuminated for 2 minutes with 405 nm LED (Lumencor, Spectra X, OR) in an inverted microscope (Nikon Instruments, Eclipse Ti-E, NY) using a 60x objective lens (Nikon Instruments, CFI Plan Apo VC 60XWI, NY). This procedure was repeated 3 to 5 times with an interval of 15-20 min. The fish were then kept in the dark until 120 hpf and imaged with a confocal microscope (Zeiss, LSM 710, Germany) (> 6 fish per time point). The expression of Kaede in the earliest born hindbrain V2a neurons, driven by the Gal4/UAS system was delayed roughly 6 hours relative to the one driven directly by the V2a promoter (data not shown).

### Two-photon calcium imaging of GCaMP6s fluorescence in hindbrain V2a neurons

The genetically encoded calcium indicator GCaMP6s (Chen et al. 2013) was expressed in hindbrain V2a neurons using the Gal4-UAS system (Tg(vsx2:Gal4; UAS:GCaMP6s)). Fish at 120-132 hpf were paralyzed with α-bungarotoxin (MilliporeSigma, 203980, MO) and then embedded in 1.6 % low-melting point agar. The agar around the tail and the nostrils was released using a fine tungsten pin. Axial motor nerve activity was recorded through fire-polished glass pipettes placed on the dorsal intermyotomal cleft on both sides. A gentle suction (∽15 mm Hg) was applied through a pneumatic device to gently break the skin. The signal was amplified (1000x) and bandpass filtered (100- 1000 Hz) through an extracellular amplifier (NPI, EXT-02B, Germany) and digitized at 6 kHz with PCIe-6363 (National Instruments, TX) using a custom C# program (available upon request). Clear motor activity was typically observed within 20 minutes. The whole hindbrain was imaged with a custom two-photon microscope equipped with a resonant scanner (Thorlabs, MPM-2PKIT) and a piezo objective scanner (PI, P-725K129, Germany) at a volume rate of 2 Hz (512 x 256, 30 slices, 7 μm z step) using Scanimage (Vidrio Technologies, VA). 940 nm, 80 MHz femtosecond laser pulses were used to excite GCaMP6s (MKS Instruments, Mai Tai HP, Deep See, MA). A gradual mechanical stimulus was delivered by gradually contacting the rostral part of the head with a fire polished glass pipette controlled by a motorized manipulator (Luigs & Neumann, Junior RE, Germany). A brief electrical stimulus (0.5-1ms in duration, 2-10 V) was delivered through a glass electrode placed on the side of the head using a stimulus isolator (Digitimer Ltd., DS-2, England). The inter-stimulus interval was from 50 sec to 2 min. The stimulus amplitude was adjusted to evoke strong motor activity consistently. Each experiment lasted between 20 and 40 min and thus contained 2400-4800 stacks.

### Analysis of two-photon calcium imaging data

Two-photon data was first corrected for mismatch between the odd and even scan lines by cross correlation if necessary and then the shift of the sample was corrected by cross correlation to a reference volume. Then GCaMP6s signal from identifiable V2a neurons were examined in relationship to fictive swim bouts. The regions of interest (ROI) for V2a positive reticulospinal neurons (Mid2i, Mid3i, RoM2, RoM3, RoV3, dorsal MiV1, and ventral MiV1/2) were drawn based on their distinct segmental distribution, dorsoventral position and soma morphology. This was confirmed further with spinal backfill (Kimmel, Powell, and Metcalfe 1982). The ROIs for caudal hindbrain V2a neurons were drawn based on their stereotypical positions as examined in the birthdating analysis: the early-born caudal neurons were displaced laterally from the rest of caudal V2a neurons while the late-born caudal neurons were located dorsally, close to the midline. Then the fluorescence time course (F(t)) for each ROI was extracted and the baseline fluorescence (F_0_) was estimated as the bottom 20^th^ percentile of the whole timeseries. Then ΔF(t)/F_0_ was calculated as follows.

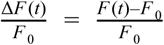

To detect weak spontaneous swimming activity reliably, the axial motor nerve activity was processed by taking a windowed standard deviation of the original signal with a 10 ms moving window as described previously (Ahrens et al. 2012). Then, a threshold for swimming activity was selected to detect the weak spontaneous swimming activity reliably from the transformed signal. In the gradual mechanical stimulus experiment, the latency of strong swimming activity from the onset of the stimulus was not consistent across trials. Thus, we used the following procedure to detect the strong bursting activity. For each swim bout, the maximum burst amplitude of the transformed signal was extracted independently for left and right channels (Fig. 3B i). Then, the distribution of the maximum burst amplitude was log-transformed to reduce skew in the distribution and fitted as a Gaussian mixture distribution with 2 components to estimate the distribution of the maximum burst amplitude that corresponds to weak spontaneous swims. This estimated probability distribution corresponding to weak spontaneous swims was used to compute a value whose probability of falling in this distribution is less than 0.1% (Fig. 4B i). The swim episodes with the maximum burst amplitude above this value were categorized as strong swims induced by the stimulus (Fig. 4B i; Fig. 3C, “Push”). The detected episodes matched with the slow and strong bursting activity (as assessed by eye). In the electrical pulse stimulus experiment, fast swim episodes were elicited reliably by the stimulus and were categorized as shock-induced fast swims (Fig. 3C, “Shock”). The remaining spontaneous swim episodes were categorized as spontaneous swims (Fig. 3C, “Sponta swim”). To derive the ΔF/F0 response related to these swim events (Fig. 3C), ΔF/F0 signal was aligned at the onsets of a given swim type and averaged over trials for each cell. Only the fictive motor signal ipsilateral to the cell being examined was considered for the following reasons: 1) hindbrain V2a neurons are primarily ipsilaterally projecting neurons (Kinkhabwala et al. 2011; Cepeda-Nieto, Pfaff, and Varela-Echavarría 2005). 2) most of the neurons related to the strong swims responded only when there is a strong axial motor activity on the ipsilateral side (Fig. 3A, 3B). The peak of the mean ΔF/F0 response was detected for each neuron and pooled for a given cell type to test if they showed activity consistently during a given swim type (Fig. 3C, top row). To test if swim types had effects on the activity of a given cell type, an one-way ANOVA with a factor for swim types was used. All possible comparisons were tested with Tukey’s multiple comparison test (Fig. 3C, bottom row).

To map the hindbrain V2a neurons recruited during these swim bouts in an unbiased way, voxel-level regression analysis was done with regressors representing the distinct types of swim bouts. Each regressor was constructed by convolving the corresponding fictive motor signal (as defined above) with a GCaMP6s impulse response function modeled as the rise and decay exponentials (0.5 s rise and 2 s decay) and then standardized for a given experimental session by subtracting its mean from its values and then dividing these new values by its standard deviation (Fig. 4A ii, 4B ii). Voxel time course (Y) was fitted with these standardized regressors (X) using the following linear model.

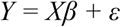

The standardized coefficient (β) was estimated by ordinary least square and then *T* value for each voxel was calculated based on standardized coefficient (β) and residual (ε). Correction for multiple comparisons was done with the false discovery rate (FDR) (Miri, Daie, Burdine, et al. 2011). The threshold for *T* maps was set at *P_FDR_* < 0.05.

### Spinal imaging of descending fibers from hindbrain V2a neurons

The photoconvertible protein, Kaede, and GFP fused to synaptophysin were expressed in V2a neurons using the Gal4/UAS system (Tg(vsx2:Gal4;UAS:Kaede;UAS:synaptophysin-EGFP)). Hindbrain V2a neurons were photoconverted with the same set up used for optical backfill but with a shorter total exposure time (1 round of 1 min exposure). The photoconversions were done at 42 and 96 hpf to visualize the spinal projections of hindbrain V2a neurons that existed at a given developmental time point. The fish were then kept in the box that filtered out UV until the imaging at 120 hpf. A series of confocal images were taken in the hindbrain and the spinal cord from the dorsal side (Zeiss, LSM710, Germany). In a separate set of experiments, hindbrain V2a neurons were photoconverted at 42 hpf and spinal motoneurons in the rostral spinal cord were labeled by the injection of a far-red dye (Thermo Fisher Scientific, Alexa Fluor 680 dextran, MW: 10,000, MA) in the axial muscles at 108 hpf. In yet another set of experiments, spinal interneurons in the rostral spinal cord were labeled instead by the injection of the far red dye in the caudal spinal cord (28^th^ to 30^th^ muscle segment) in a manner described previously (Hale, Ritter, and Fetcho 2001).

### Analysis of distribution of converted Kaede in the hindbrain and spinal neuropil

Volumetric images of V2a neurons labeled with converted and unconverted Kaede in the hindbrain and the spinal cord were 3D-rendered in Imaris (Bitplane, Switzerland). Then maximum intensity projection views from the dorsal side were created in the area of interest. The views were carefully oriented based on anatomical landmarks such as axons from V2a neurons to minimize the error from tilt of the 3D volume. Then the intensity distribution in the neuropil along the mediolateral axis in the projection images was calculated in Fiji (Fiji is just Image J) from a line profile extending from the lateral surface of the cell bodies of the most laterally located V2a neurons to the lateral surface of the neuropil from V2a neurons. The line thickness was set to 20 μm in Fiji to average the intensity profiles over 20 μm in the rostrocaudal direction. The intensity of converted Kaede and the mediolateral position were normalized in each fish and the average intensity profiles and their standard errors were plotted as a function of the normalized mediolateral position in the neuropil.

### Hindbrain whole-cell recordings

Patch-clamp recordings of hindbrain neurons were performed with a procedure similar to the one described previously (Kimura et al. 2013), but with some modifications. Five-day-old fish were paralyzed with α-bungarotoxin (MilliporeSigma, 203980, MO) that was dissolved in system water (1 mg/ml). After successful paralysis, larvae were anesthetized with MS-222 and then secured with etched tungsten wires (pins) through the notochord to a Sylgard-coated glass-bottom dish containing extracellular solution (134 mM NaCl, 2.9 mM KCl, 1.2 mM MgCl2, 2.1 mM CaCl2, 10 mM HEPES, and 10 mM glucose, adjusted to pH 7.8 with NaOH). Then, the head was rotated and secured ventral side up with tungsten pins placed through the ears and the rostral part of the jaw (Fig. 7A ii). The ventral surface of the hindbrain was carefully exposed by removing the notochord using an tungsten pin and fine forceps. The skin of the middle region of the body was removed with a pair of forceps to gain access to the peripheral nerves from spinal motoneurons. Locations of motor nerve recordings were from the 10^th^ to 15^th^ muscle segment.

Neurons were targeted based on fluorescence and scanned Dodt gradient contrast images acquired with a custom two-photon microscope. 40x (Nikon Instruments, CFI Apo 40XW NIR, NY) or 25x (Leica, HCX IRAPO L 25x/ 0.95 W, Germany) objective lens was used. Descending hindbrain neurons were identified by the fluorescence signal from Tg(vsx2:EGFP), Tg(vsx2:Kaede) or spinal backfill (see above). MiV1 neurons were located in rhomobomere 4, medial and ventral to Mauthner cells which can be easily identified by Dodt gradient contrast. The subclass of MiV1 neurons was targeted based on photoconversion of Kaede (see above) and its stereotypical position.

Whole-cell recordings were established using the standard procedure and the signals were recorded using EPC 10 Quadro amplifier (HEKA Electronik, Germany) and PatchMaster (HEKA Electronik). The resistance of the electrode was 8 to 12 MOhm. The intracellular solution contained (in mM) 125 mM K-gluconate, 2.5 mM MgCl2, 10 mM EGTA, 10 mM HEPES and 4 mM Na2ATP adjusted to pH 7.3 with KOH. A junction potential using this intracellular and extracellular solution (see above) has been calculated at 16 mV, which would result in a shift of measured potentials 16 mV in the negative direction. Because this would not affect our conclusions, we did not correct for it. The solution also contained 0.01% of Alexa Fluor 568 hydrazide or Alexa Fluor 647 hydrazide (Thermo Fisher Scientific). Z stacks were acquired with a custom two-photon microscope to confirm morphology and locations of the recorded neurons. Input resistances were calculated from an average of 5 hyperpolarizing square current pulses between 20 and 80 pA. This range of current injection produces a linear current-voltage response in all the MiV1 neurons examined.

Electrophysiological analysis of recruitment of MiV1 neurons was performed in a manner similar to the previous studies (McLean et al. 2007, 2008). Fast fictive swimming was induced by a brief electrical stimulus ( < 1 ms in duration at 1-10 V) delivered via a pair of tungsten electrodes (A-M systems, WA) placed near the tail. Slow swimming often occurs spontaneously but is also induced by flashes of light. Cycle frequencies of axial motor activity were computed by taking the reciprocals of the time interval between each burst and the next one. The spiking and subthreshold activity that precede each cycle were extracted and their relationship to cycle frequency was examined as follows. Spikes were detected based on the threshold determined for each cell. Subthreshold activity was examined based on the lowpass-filtered (60Hz) voltage trace, which effectively isolated rhythmic subthreshold activity during swimming. The maximum depolarization for each cycle was calculated by subtracting its peak from the baseline which was defined as the minimum voltage in the 200-ms time window before the swim onset. In order to statistically test if each age group changes spiking and subthreshold activity as a function of cycle frequency, each cycle was classified as fast (> 35 Hz) or slow (< 35 Hz) cycle and fitted with the following linear mixed models with a random effect for cell using the *nlme* package in R (https://cran.r-project.org/web/packages/nlme/index.html). Number of spikes was log-transformed to satisfy the assumptions of the linear model. The effect of cycle speed was tested by comparing the activity for each age group with correction for multiple comparisons using the *multcomp* package in R (https://cran.r-project.org/web/packages/multcomp/index.html).

*log (number of spikes)* ∼ *cycle speed* ^*^ *age*

*membrane depolarization* ∼ *cycle speed* ^*^ *age*

### Single cell analysis of spinal projection

Electroporation of Alexa Fluor 647 dextran dye (MW 10,000; Thermo Fisher Scientific) into single neurons was performed as described previously (Koyama et al. 2011). MiV1 neurons were targeted based on GFP expression in Tg(vsx2:GFP) and/or spinal backfill with Texas Red dextran dye (MW 10,000; Thermo Fisher Scientific) along with scanned Dodt gradient contrast under a custom two-photon microscope at 5 days old. One day after electroporation, the hindbrain was imaged with a confocal microscope (Zeiss, LSM 710) from the dorsal side to confirm the dorsoventral position of the electroporated MiV1 neurons. Multiple z stacks from the side were also imaged from hindbrain to the caudal end of spinal cord to cover the entire spinal projection. Stacks were then stitched together with stitching plugin available through Fiji (Preibisch, Saalfeld, and Tomancak 2009). When spinal backfill was used to visualize descending neurons, the spinal segments caudal to the injection site (21^st^ to 23^rd^ muscle segment) were excluded from imaging. Stitched stacks were 3D rendered in Imaris (Bitplane, Switzerland). To visualize the spinal projection in coronal view, axonal arborization was reconstructed with neuTube (http://www.neutracing.com/) and then visualized in Imaris. Three to four muscle segments were used to render coronal views in the rostral spinal cord (6^th^ to 9^th^ muscle segment) and in the middle spinal cord (14^th^ to 17^th^ muscle segment). All the cells examined in the study had their main axon in the ventral spinal cord and had collaterals innervating the dorsal spinal cord. The dorsal extent of the axonal arborization was quantified relative to the total dorsoventral extent of spinal cord as described previously (McLean et al. 2007). To statistically examine if the extent of spinal arborization depends on hindbrain cell type and the position in the spinal cord, the dorsal extent of arborization was fitted using the following linear mixed effects model with a random effect for cell using *nlme* package in R.

*Dorsal extent of axon collaterals* ∼ *hindbrain neuron types* ^*^ *position in the spinal cord* + (1|*Cell ID*)

Putative presynaptic terminals were identified based on their characteristic puncta-like structure. Then their apposition to fluorescently labeled spinal neurons was examined. Primary motoneurons were identified using Tg(mnx1:TagRFPT). Secondary motoneurons were identified using TgBAC(islet1:EGFP). Spinal V2a neurons (also called circumferential ipsilateral descending neuron (CiD)) were identified using TgBAC(vsx2:EGFP). Multipolar commissural neurons (MCoD) were identified by spinal backfill from the caudal spinal cord (see above). To statistically examine if hindbrain cell type explained the variance in their spinal innervation patterns, the presence of innervation to the spinal neurons was fitted with the following generalized linear mixed effects model with a random effect for cell and a logit link function for a binomial distribution using *lme4* package in R (https://cran.r-project.org/web/packages/lme4/index.html).

*Presence of innervation* ∼ *hindbrain neuron types* ^*^ *spinal neuron types* + (1| *Cell ID*)

### Paired whole-cell recordings of hindbrain and spinal cord neurons

Whole-cell recordings from hindbrain neurons were done as described above. The spinal cord was also exposed by removing the muscles overlying the cell of interest using the standard procedure described previously (Drapeau et al., 1999). Dorsal and ventral MiV1 neurons were targeted as described above. Mid3i was identified based on the GFP signal from Tg(vsx2:EGFP), its stereotypical segmental location (rhombomere 6) and morphology (laterally elongated soma and its position relative to nearby reticulospinal neurons in the same segment). Mauthner cell was identified based on its segmental position (rhombomere 4) and its large soma which is clearly visible under Dodt gradient contrast. Spinal motoneurons and interneurons (MCoDs) were targeted with Dodt gradient contrast and/or fluorescence either from backfill (see above) or from transgenic lines (Tg(mnx1:TagRPFT) for motoneurons; Tg(Dbx1b:Cre, vglut2a:lRl-GFP) for MCoDs). Primary motoneurons are dorsally located large cells close to the lateral surface of the spinal cord. MCoDs are ventrally located smaller cells but also close to the lateral surface of the spinal cord. PMNs were sampled at muscle segment 17 to 19. MCoDs were sampled at muscle segment 11 to 13. To examine if a given connection is monosynaptic, extracellular solution with a high concentration of divalent cations (4x Mg2+ and 4x Ca2+) was perfused to increase the firing threshold of potential relay neurons. Only the connection between Mauthner cell and MCoD was almost completely abolished, which is also consistent with its long latency. Thus this connection was excluded from subsequent analysis. To examine the composition of electrical and chemical synapses for a given connection, chemical synapses were blocked by a cocktail of D-AP5 (Tocris, 0106, MN) and NBQX (Tocris, 0373, MN) for glutamatergic synapses and by mecamylamine (Tocris, 2843, MN) for cholinergic synapses.

To statistically examine if hindbrain and spinal cell types had significant effects on the presence of monosynaptic connections, the presence of monosynaptic connection was fitted with the following generalized linear model with a logit link function for a binomial distribution using *lme4* package in R.

*Presence of monosynaptic connection* ∼ *hindbrain neuron types* ^*^ *spinal neuron types*

The differences in conduction velocity, postsynaptic potential (PSP) amplitude and PSP half decay time across connected pairs of the monosynaptic responses were statistically examined with Kruskal-Wallis test. Post hoc comparisons were done with Dunn’s multiple comparison test.

### Targeted two-photon laser ablation

We ablated two groups of V2a neurons based on the developmental and recruitment analyses we performed (Fig. 2-4). The first group consisted of the early-born V2a neurons that were recruited during strong swims. Procedurally, they were defined as hindbrain neurons that contained photoconverted Kaede at 60 hpf after photoconversion at 24 hpf in Tg[vsx2:Kaede] (Fig. 10B i). The second group consisted of the late-born V2a neurons in the caudal hindbrain that were recruited during spontaneous weak swims. Procedurally, they were defined as neurons that did not contain photoconverted Kaede at 60 hpf after photoconversion at 42 hpf and met the following spatial criteria (Fig. 10B iv). First, their somata were caudal to rhombomere 6 where the early-born neurons were located in dorsal hindbrain (Kinkhabwala et al. 2011) Second, they were dorsal to the early-born neurons in the caudal hindbrain. We chose these definitions to capture two distinct and minimal sets of neurons that we predicted would cause distinct locomotion phenotypes based on their developmental and recruitment profiles. We also matched the time of ablation in the two groups.

For every fish that was randomly selected for ablation from the pool of fish that had been photoconverted, one or two more fish were chosen in the same way to serve as a negative control(s). The fish to be ablated was anesthetized in MS-222 for 2 minutes before being mounted dorsal-side up in 1.6% low melting point agarose and placed under our custom two-photon microscope where the ablations took place. Both the fish in which the ablations were carried out and the fish chosen as control were kept under anesthesia for the entire duration of the ablations. Before proceeding to ablations, we first imaged hindbrain V2a neurons labeled with unconverted and converted Kaede using 1020 nm, 80 MHz femtosecond laser pulses (MKS Instruments, Insight, Deep See, MA) with a 25x objective lens (Leica, HCX IRAPO L 25x/ 0.95 W, Germany). Then we proceeded to ablations starting from the most ventral target cells to increasingly more dorsal cells to minimize light scattering that could result from preceding ablations. For a given depth, each target cell was exposed to femtosecond laser pulses for 5 ms with spiral scans centered on its soma (outer diameter, 3um) before targeting the next target after a minimum 5 second interval using Scanimage (Vidrio Technologies, VA). We used 920 nm or 1020 nm laser (196 mW or 142 mW after objective lens respectively). This procedure was repeated up to ten times until the cells at a given depth were deemed ablated based on visual inspection. After ablating all the cells of interest, we imaged hindbrain V2a neurons again to assess the quality of ablation. Then, the fish was gently removed from the embedding agarose and both this fish and the control fish were housed singly in separate dishes until later use on the day of behavior imaging, which was at 5 dpf. To examine the potential recovery of target cells after the ablation, a subset of the ablated fish and the control fish was selected for imaging at 4 dpf. From the time of photoconversion till the day of behavior, the dishes containing the fish were enclosed in our UV-opaque box to prevent further conversion of Kaede.

### Behavioral assays

At 5 dpf, ablated and control fish were transferred singly and in random sequence to a 50 mm diameter circular arena where they were allowed to swim freely while being imaged from the top with a high speed camera (Miktrotron, MC-1362, Germany). To confine the movements of the fish to the yaw (x-y) plane, the depth of the water in the arena was limited to 2 mm. Before the start of imaging, each fish was allowed to acclimate to the arena for a minimum of 20 minutes before their behavior was imaged. If even after 20 minutes the fish spent a majority of the time being stationary or swimming very close to the edge of the arena (as they tend to do soon after being transferred into the arena from their home dish), then the wait period was increased until the fish started showing frequent swims in trajectories that took them far from the arena edge. Following this acclimation period, fish were first imaged at 300 frames per second (fps) for a total 5 minutes while they swam spontaneously. After this 5 minute period, fish were imaged continuously while they were presented with a series of alternating vibration and dark flash stimuli - although in this paper we did not use data from dark flash responses - at 1 minute intervals. A time period of 100 ms before and 1400 ms after each stimulus were imaged at 500 fps, whereas the intervening 1 minute periods between stimuli were imaged at 30 fps. The vibration stimulus was delivered via a sound speaker in contact with the acrylic base plate upon which the fish arena was placed. The sound stimulus consisted of two cycles of 500 Hz sine wave at (peak-to-peak acceleration ≈ 100 *ms^-2^*). For the dark flash stimulus, the ambient light was turned off for 300 ms.

Images collected during the behavior sessions were first sorted by frame rate. For images collected at 30 fps we only tracked the fish’s head position, whereas for images collected at 300- and 500 fps, we estimated the fish’s head orientation and body curvature as well. To extract the fish’s head position, we first computed a background image and subtracted this from all images to remove background from them. The background image was computed by averaging a set of 1000 temporally uniformly spaced images from the set of all images. After background subtraction, for convenience, the pixel values were multiplied by −1 so that the fish’s eyes would go from being the darkest to the brightest spots in the image. The images were then smoothed by convolving each image with a 1 mm wide 2D Gaussian kernel. The convolved images were then segmented using an automatically estimated threshold (multithresh function in MATLAB) to isolate within each image a blob (contiguous set of pixels) of the brightest pixels that included the fish’s eyes as well as the swim bladder. The centroid of these isolated pixels was used as the fish’s head position. The centroid obtained in this manner was reliably located between the eyes and the swim bladder in all image frames and for frames in which the fish did not move, the centroid typically shifted by less than 0.17 mm (≈ 3 pixels) between frames (confidence interval of 95%).

For quantifying the fish’s tail curvature we needed to reliably isolate as many pixels on the fish’s body as possible so that the movements of the tail could be accurately tracked during swimming. For this we used a custom MATLAB script that used an iterative procedure to obtain a threshold for image binarization and isolation of fish pixels. The flowchart briefly describing this procedure is shown below. Our algorithm relied on the assumption that a single threshold automatically estimated by MATLAB’s multithresh function would reliably isolate only a subset of fish pixels, but that less than two-thirds of the actual number of fish pixels would remain to be isolated still following this segmentation procedure. We arrived at this assumption empirically when trying to devise an algorithm that could reliably detect fish pixels from within our background subtracted images. By using the procedure described above, we were able to isolate fish pixels reasonably well in our image datasets.

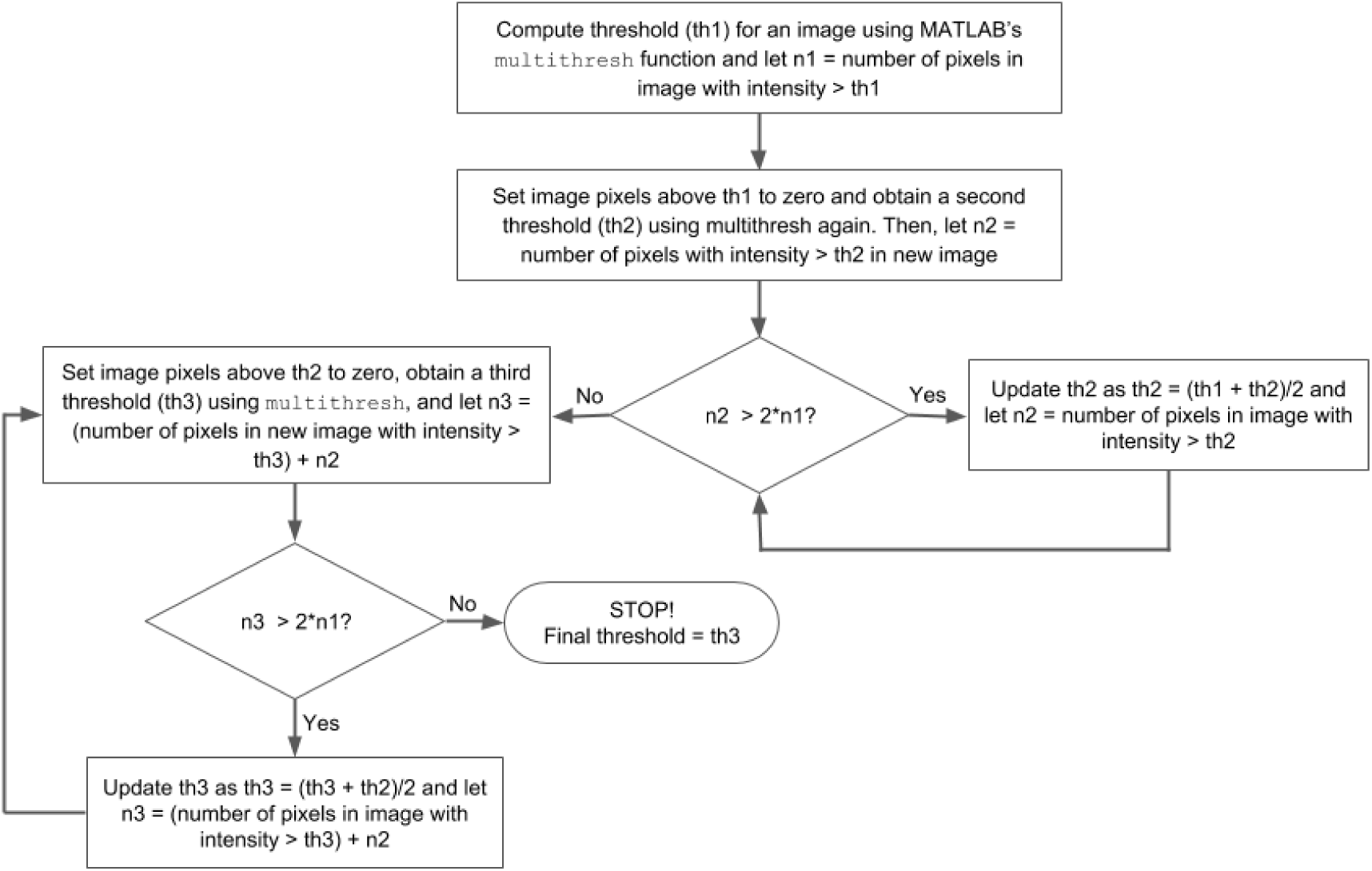

Following the detection of fish pixels, the images were binarized by setting fish pixels to 1 and the rest of the pixels to 0. The MATLAB function bwmorph was then used on the binary images to “thin” the fish pixels down to a set of connected pixels that spanned the length of the fish and coarsely bisected it into two lateral halves. To transform these “midline” pixels into a smoother curve that better tracked the spine of the fish, we followed the procedure described in (Huang et al. 2013), wherein the integer-valued image coordinates of the midline pixels (*x_i_*, *y_i_*) were weighted by the intensities of the surrounding pixels to obtain a new decimal-valued set of pixels 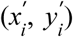 as follows:

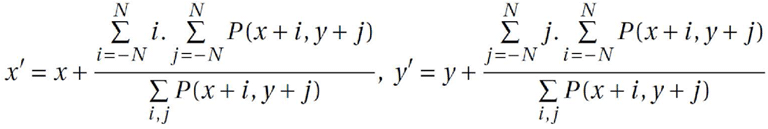

Where *P* represents the pixel intensities and *N* is the range of the surrounding pixels, which in this case is 1. The curve obtained by this procedure was then interpolated using cubic splines to obtain a final smooth curve (*C*) of 50 points.

### Quantification of free swimming behaviors

To characterize swimming we looked at changes in the body posture and head orientation over time. We used the midline curve (*C*) obtained using the procedure described above to characterize body posture. First, we computed tangent vectors 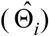 along the points on *C* as follows

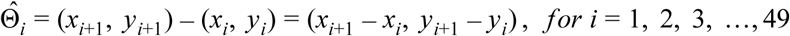

Next, each of these tangent vectors 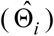 was transformed into a point on the complex plane (*z_i_*) such that

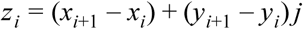

where *j* is the imaginary number 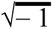. Then, local curvatures (κ_i_) along the points on *C* were calculated as follows

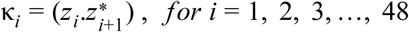

where for a given a complex number *z*, *Arg*(*z*), can be computed using the MATLAB function angle. In the above equation, 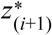 indicates the complex conjugate of *z*_*i*+1_. Finally, we spline-interpolated the set of local curvatures obtained by the above procedure to obtain a final set of 50 local curvature values 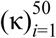 along the curve *C*. To quantify the total amount by which a fish bent its body for a given time point *t* we simply added all local curvatures along the midline curve *C*(*t*) associated with that time point. Thus, the total curvature (K) for a given time point was given by

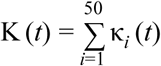

To obtain the head orientation, the first 10 points (the 10 points closest to the head centroid) of the midline curve *C* were sampled and a straight line (*L*) was fit to those points. The head orientation (*φ_H_*) was then estimated as the absolute angle of the vector extending from the point on *L* farthest from the head centroid to the point on *L* closest to the head centroid.

To identify distinct swim episodes, we used the total curvature time series, K(*t*), after low pass filtering it at 70 Hz to remove high frequency noise. We then used a custom MATLAB script to identify the times corresponding to the onset and offset of each swim episode. Briefly, the absolute function of the filtered curvature timeseries (| K(*t*)|) was convolved with a 200 ms gaussian kernel and in the first step the onsets of swim episodes were identified as the points where the transformed timeseries ( *S* (*t*)) crossed and stayed above a threshold of 4° for at least 50 ms. Then, for each onset a corresponding offset was identified as the first point in *S*(*t*) after this onset where the signal returns and stays below the threshold for at least 50 ms. Then, in a second step to improve on the loss of temporal resolution resulting from the convolution of K(*t*) to get *S*(*t*), we updated each onset by replacing it with the time of the peak in the second derivative of *S*(*t*) that occurred immediately before this onset. Bend amplitudes and bend periods within a swim episode were computed after detecting peaks within the curvature time series as follows

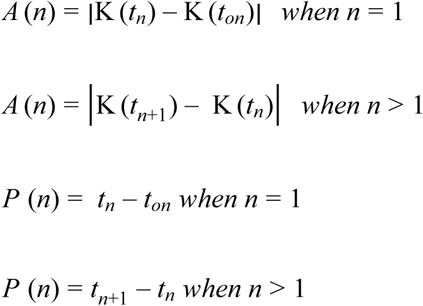

where *n* corresponds to bend number, *A* (*n*) and *P* (*n*) are the *n^th^* bend amplitude and period respectively, *t_n_* is the time at the *n^th^* peak, and *t_on_* is the time of swim onset. Peak angular velocities within a swim episode were computed in a similar manner from the angular velocity time series ω(*t*), which is the temporal derivative of the timeseries *K*(*t*).

Previous studies involving ablation of other neurons had shown that deficits resulting from ablation can be specific to a particular bend within a particular type of swimming (Huang et al. 2013). Therefore, to determine what effects ablation of V2a neurons might have had on swimming we examined the amplitudes and periods of the individual bends that constituted a swim episode. To ensure that such bend-by-bend comparisons were being made between similar swim types recorded from control and ablated fish we first sorted spontaneous swim episodes into distinct subtypes. To do so, we used the amplitudes of the first bend within each swim episode because it was shown in a previous study (Burgess and Granato 2007) that this single metric could be used to effectively categorize spontaneous swims into at least two distinct groups. To obtain the decision boundaries for sorting spontaneous swimming episodes from our dataset into distinct swim categories we extracted the first bend amplitude from each episode and fit a Gaussian Mixture (GM) model to the data using the scripts available in the scikit-learn library (http://scikit-leam.org/stable) in Python. For training the GM model we used data from 15 fish (6056 swim episodes) in which V2a neurons had not been photoconverted (Fig 10A, iv-v) because photoconversion could have potentially altered swimming characteristics. Our model incorporated three Gaussian components because this resulted in the lowest value for the Akaike Information Criterion (AIC) (Fig 10A, iv bottom), indicating that our dataset consisted of three distinct spontaneous swim types. Finally, we used the trained GM model on the entire spontaneous swimming dataset to assign each episode to one of the 3 categories. To visually inspect the swim episodes from each category we used a custom script to temporally align all the episodes (curvature time series) within a category in such a way as to produce the highest correlations among them, and then plotted the traces atop of each other (Fig 10A, vi). For aligning escape episodes which have highly nonstationary burst frequencies, correlations were computed using a restricted time window of 80 ms after the episode onset. As with the training of the GM model, we only used data from non-photoconverted fish to make these plots. The plots showed that the swim episodes within each category were quite similar not just with respect to the amplitudes of their first bends, which we had used to predict swim categories, but with respect to subsequent bends as well. Furthermore, the overlaid traces revealed that swim episodes within a category were very similar to each other, but swim episodes across categories were quite distinct. This confirmed that as in the study by Burgess and Granato (2007), we could use the amplitude of just the first bend of each swim episode to effectively categorize spontaneous swims into distinct subtypes.

In the case of escape swims, we treated them as belonging to a single category because they were all produced in response to the same vibration stimulus and because they are very rarely, if ever, observed in the absence of a stimulus. Thus, for comparing the individual bend amplitudes and periods between control and ablated fish, we did not sort escapes in the same way as spontaneous swims.

To determine if bend amplitudes and periods were significantly different in control and ablated fish, we carried out statistical tests using software packages implemented in R. The data was first fit using a mixed effects model (*nlme* package) in which the swim parameter of interest (bend amplitude or bend period) was treated as the response variable while the categorical variables treatment, ablation group, and swim type were treated as the explanatory variables. The variable treatment took two values that indicated if the source of a given data sample was an ablated or a control fish. The variable ablation group also assumed two values that indicated if a data point had been collected from a fish in which old (birth-time < 24 hpf) or young (42 hpf < birth-time < 60 hpf) alx neurons had been ablated. Finally, the variable swim type indicated which of the aforementioned swim categories the data point belonged to. In addition to the above explanatory variables, the identity of the fish was modeled as a random variable; the values within this last variable uniquely identified each fish that contributed to our dataset. The model also took into account interactions between the independent variables. We used this model-based approach to separately assess the effects of ablation on each of the first six bends for both bend amplitude and bend period. Significance was assessed at p < 0.001.

*bend amplitude* ∼ *treatment*^*^*ablation group* + *swim type* + (1| *fish ID*)

*bend period* ∼ *treatment*^*^*ablation group* + *swim type* + (1| *fish ID*)

## Acknowledgements

We are grateful to the present and past members of our lab, especially to Jamien Shea who supported the early phase of this work and Masashi Tanimoto who provided critical feedback to the manuscript. We are also thankful to James Fitzgerald, Tzumin Lee, Misha Ahrens and Erik Snapp for providing critical feedback and to the members of Misha Ahrens’ lab for their feedback during joint lab meetings. We thank all the members of the aquarium facility at Janelia for providing excellent care to the fish used in this study. We greatly appreciate the expertise and equipment provided by Janelia Experimental Technologies, especially by Vasily Goncharov and Christopher McRaven. Finally, we thank the scientific and administrative community at Janelia for creating an environment that allows for the focused pursuit of fundamental scientific questions.

---

**Figure 2 - figure supplement 1.**
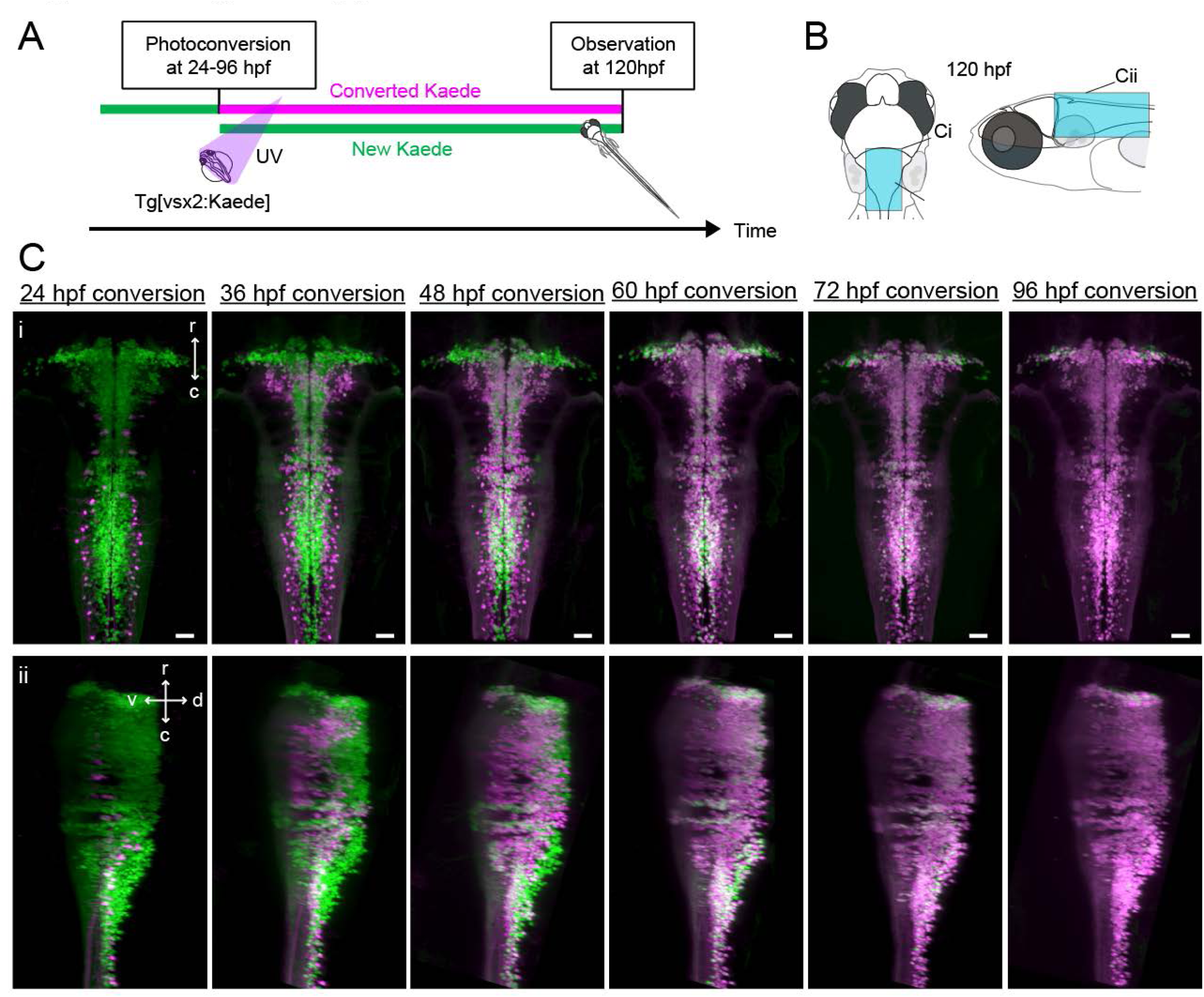
Birthdating of hindbrain V2a neurons. **A.** Experimental procedure. **B.** Regions displayed (cyan patch). **C.** Top-down (i) and side (ii) views of hindbrain V2a neurons at 120 hpf showing Kaede photoconverted at a specific time point (24 hpf – 96 hpf). Magenta, converted Kaede; green, unconverted Kaede; r, rostral; c, caudal; d, dorsal; v, ventral; Scale bars, 30 um.

**Figure 5 - figure supplement 1.**
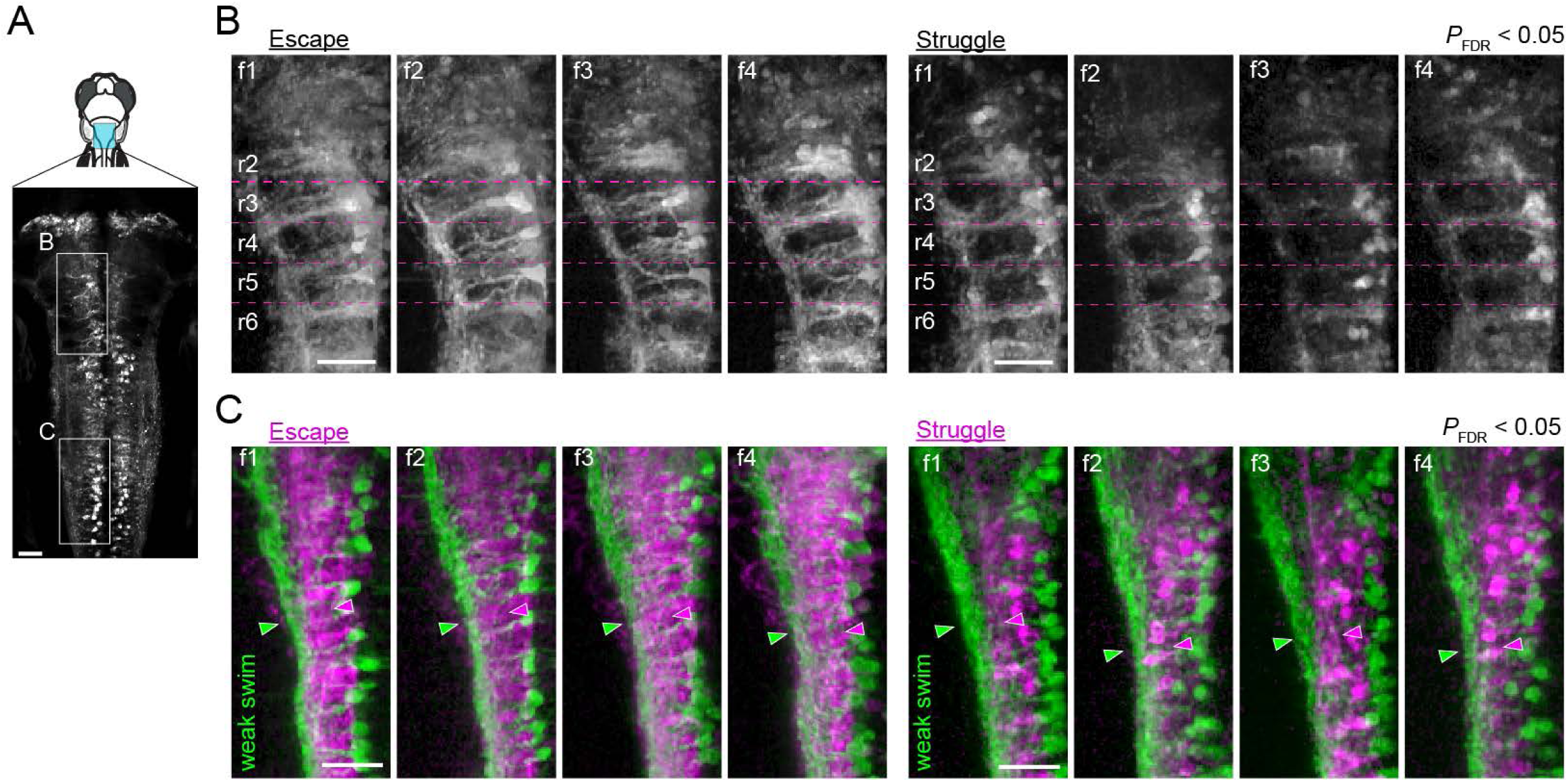
Activity maps from multiple subjects in the rostral and the caudal hindbrain. **A.** Regions displayed (cyan patch). **B.** Activity maps in the rostral hindbrain from multiple fish (f1 to f4). Dotted magenta lines indicate the boundaries between hindbrain segments (rhombomere 2 (r2) to rhombomere 6 (r6)). More V2a neurons are recruited consistently for putative escape. **C.** Activity maps in the caudal hindbrain from multiple subjects (f1 to f4). Arrowheads highlight the functional segregation of neuropil. Scale bars, 30 um.

**Figure 7 - figure supplement 1.**
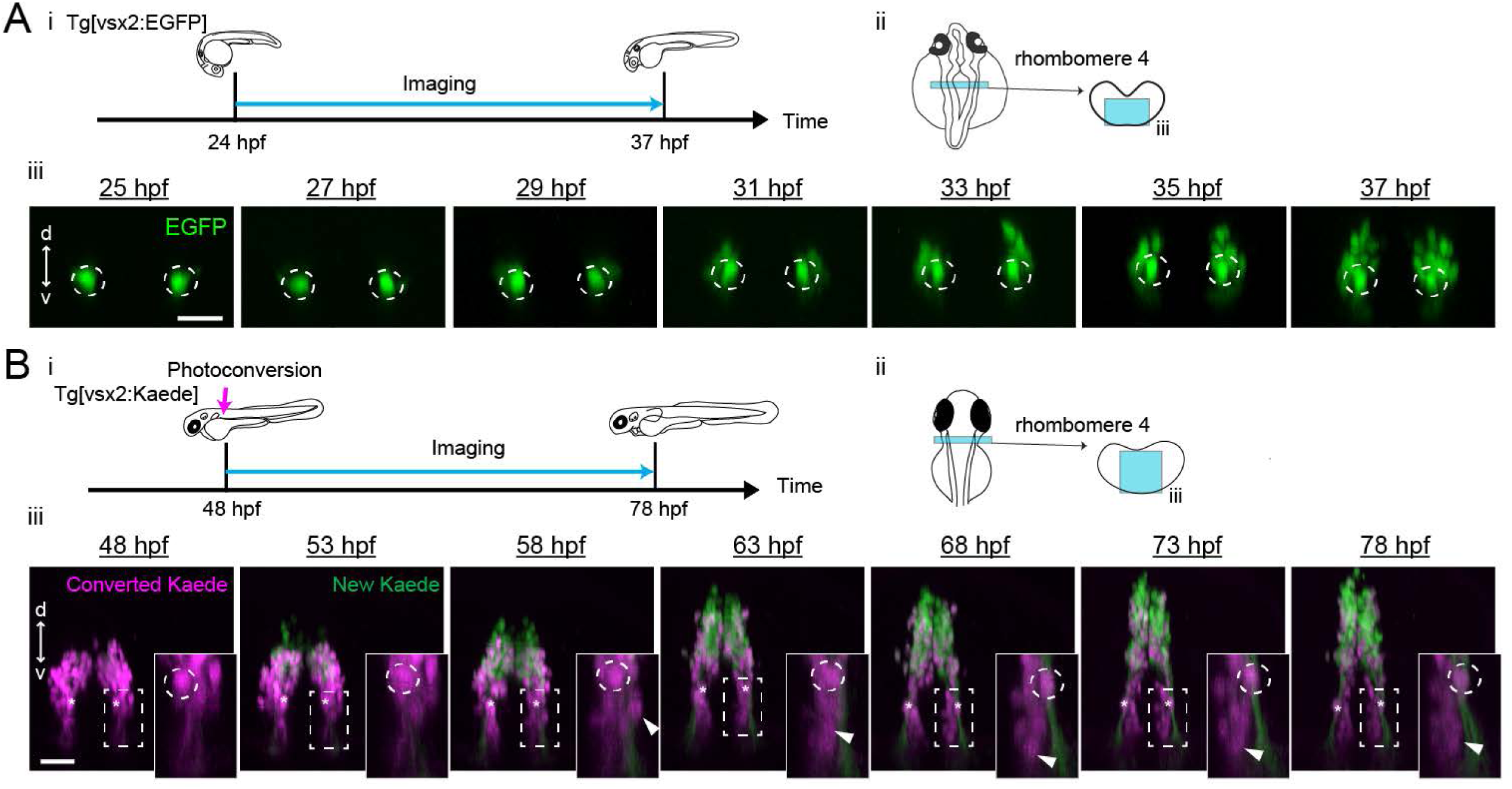
Emergence and migration of hindbrain V2a neurons in rhombomere 4. **A.** Emergence of V2a neurons in rhombomere 4. (i) Experimental procedure (ii) Region displayed (cyan patch) (iii) Time lapse coronal views of hindbrain V2a neurons in rhombomere 4 from 25 hpf to 37 hpf. Dotted circles indicate the locations of the earliest born neurons in rhombomere 4 (dorsal MiV1). d, dorsal; v, ventral. Scale bars, 30 um. **B.** Migration of V2a neurons in rhombomere 4. (i) Experimental procedure (ii) Region displayed (cyan patch). (iii) Time lapse coronal views of hindbrain V2a neurons expressing Kaede from 48 hpf to 78 hpf after photoconversion at 48 hpf. Asterisks indicate the locations of the earliest born neurons in rhombomere 4 (dorsal MiV1) tracked throughout the experiment. Dotted rectangles show the location of magnified images in insets. (inset) Dotted circle indicates dorsal MiV1. White arrowhead indicates the V2a neurons migrating ventrally past dorsal MiV1. d, dorsal; v, ventral. Scale bars, 30 um.

**Figure 10 - figure supplement 1.**
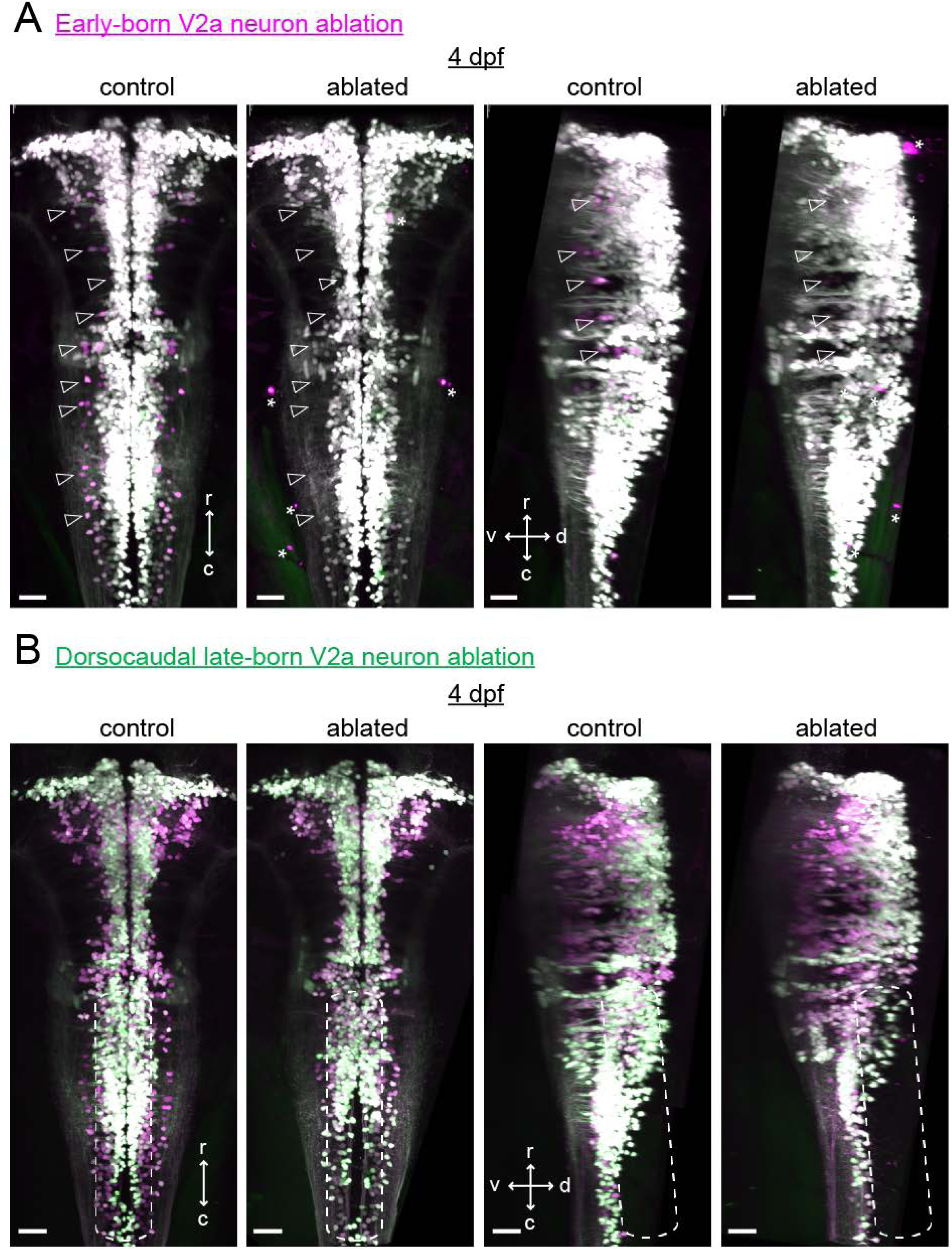
Confirmation of hindbrain V2a neuron ablations at 4 dpf. **A.** Maximum intensity projections of hindbrain V2a neurons expressing Kaede from a fish in which early-born neurons were ablated (see Figure 11B i-ii) as well as from a control fish at 4 dpf (left, dorsal view; right, side view; magenta, photoconverted Kaede; green, unconverted Kaede). Contrast of magenta channel was enhanced to improve the visibility of early-born neurons, which made late-born neurons appear white. Open arrowheads indicate locations of early-born neurons (magenta). Note the lack of magenta cells in the ablated fish. Asterisks mark the bright structures in the red channel located outside the brain. r, rostral; c, caudal; d, dorsal; v ventral; scale bars, 30 um. **B.** Maximum intensity projections of hindbrain V2a neurons from a fish in which dorsocaudal late-born neurons were ablated (see Figure 11B iii-iv) as well as a control fish at 4 dpf (left, dorsal view; right, side view; magenta, photoconverted Kaede; green, unconverted Kaede). Rounded dotted rectangle in each image indicates the location of neurons that were targeted in the ablated fish. r, rostral; c, caudal; d, dorsal; v ventral; scale bars, 30 um.

**Figure 10 - figure supplement 2.**
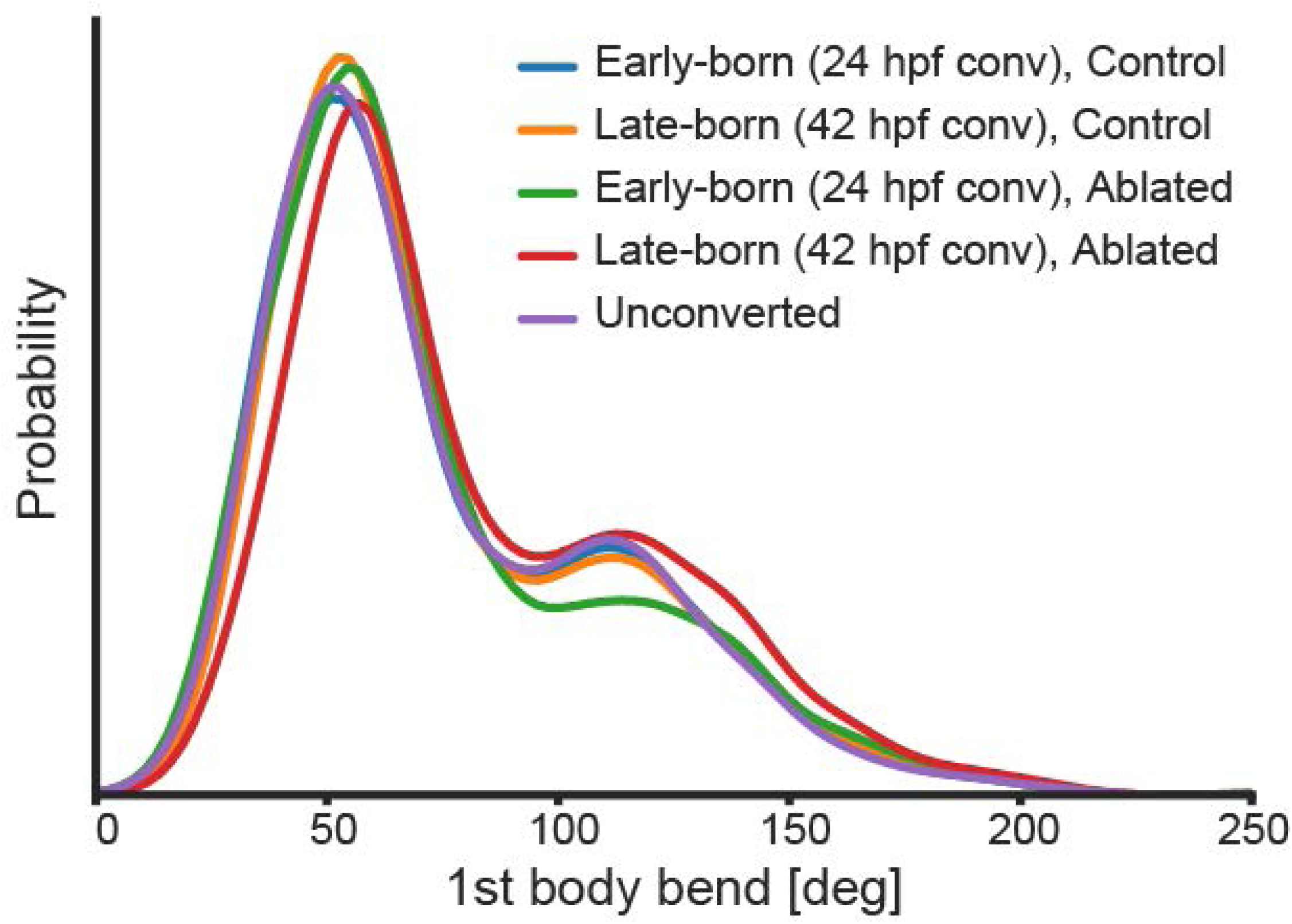
Probability distributions of the first body bend amplitude of spontaneous swim events in the ablated and control groups. Blue, the early-born neurons ablation group (24 hpf phoconversion, 10 fish, 1156 episodes); orange, the late-born neurons ablation group (42 hpf photoconversion, 10 fish, 1067 episodes); green, the early-born neurons control group (24 hpf photoconversion, 8 fish, 2195 episodes); red, the late-born neurons control group (42 hpf photoconversion, 8 fish, 1475 episodes); purple, unconverted control group (15 fish, 1175 episodes).

**Figure 10 - figure supplement 3.**
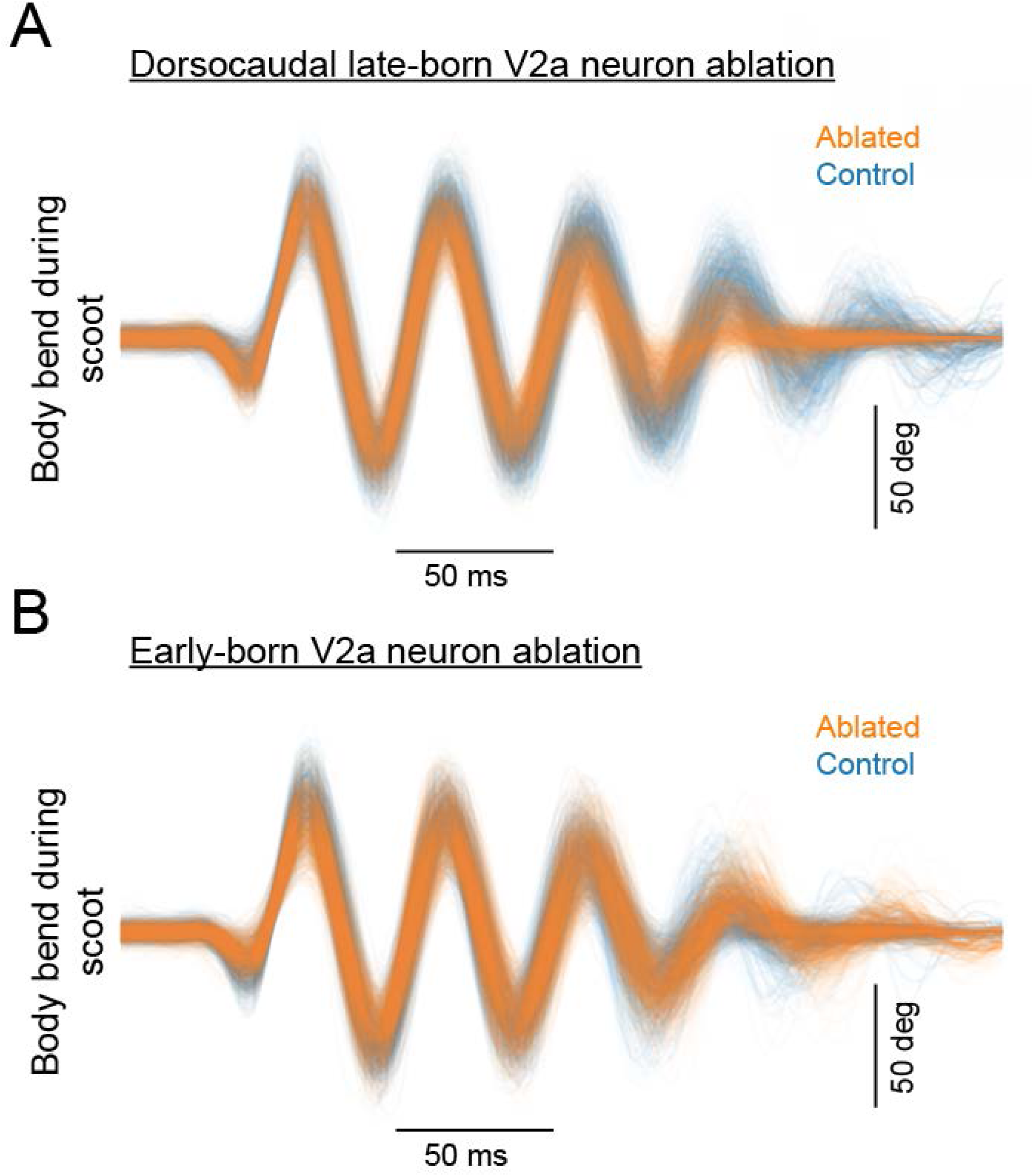
Overlaid traces of body bend during scoot in the ablated and control groups. **A.** Overlaid traces of body bend during scoot in the dorsocaudal late-born V2a neuron ablation group and the corresponding control group. Orange, ablated (10 fish, 853 episodes); Blue, control (8 fish, 901 episodes). **B.** Overlaid traces of body bend during scoot in the early-born V2a neuron ablation group and the corresponding control group.Orange, ablated (10 fish, 815 episodes); Blue, control (8 fish, 791 episodes).

**Supplementary video 1. Ca^2+^ activity of hindbrain V2a neurons during fast and slow swimming episodes.**

Top-down view of GCaMP6s-labeled hindbrain V2a neurons showing activity correlated with distinct locomotor patterns. Dorsal side of the brain is at the top of movie frames. Axial motor activity is encoded with a sound signal. The time of the electrical stimulus delivered to the right side is indicated by a red square patch in the lower right side. The movie is sped up 7.5 times.

**Supplementary video 2. Migration of hindbrain V2a neurons in rhombomere**

Coronal view of hindbrain V2a neurons expressing Kaede in rhombomere 4 imaged from 48 hpf to 84 hpf. Kaede photoconverted at 48 hpf is shown in magenta and unconverted Kaede expressed afterwards is shown in green. The earliest-born V2a neurons in rhombomere 4 (dorsal MiV1) are indicated by two gray ellipsoids, one on each side. Note the magenta cells migrating past dorsal MiV1.

